# Super-resolved imaging of mRNA ultrastructure in cells

**DOI:** 10.64898/2026.01.23.701081

**Authors:** A. Passera, P. Welzl, A. Gomez-Segalas, C. Plaschka, F. Balzarotti

**Affiliations:** Research Institute of Molecular Pathology (IMP), Vienna BioCenter (VBC); Campus-Vienna-Biocenter 1, 1030 Vienna, Austria; Vienna BioCenter PhD Program, Doctoral School of the University of Vienna and Medical University of Vienna; A-1030, Vienna, Austria

## Abstract

Messenger RNA (mRNA) is a central polymer of gene expression, whose ultrastructure and defining regulatory rules remain unclear. To visualize the ultrastructure of mRNA, we here develop Combi-PAINT, a powerful and generalizable method for combinatorial super-resolved DNA-PAINT multiplexing, which we combine with high-efficiency RNA labelling and MINFLUX microscopy. This approach enables the nanometer-precision tracing of mRNA in three dimensions in cells. We use Combi-PAINT to visualize multiple distinct mRNA species inside the human cell nucleus and cytoplasm, revealing their quantitative ultrastructures and transcript-specific molecular patterns. By bridging sequence specificity with nanometer resolution, we provide a new lens for studying the cellular life of mRNA.

## Introduction

The three-dimensional organization of messenger RNA (mRNA) is critical for cellular function. mRNAs adopt secondary and tertiary structures and assemble with RNA-binding proteins (RBPs) into messenger ribonucleoprotein particles (mRNPs) (*1–5*), which govern fundamental processes such as RNA processing, packaging, diffusion, nuclear export, translation, and decay. Sequencing-based approaches can map RNA secondary structures (*6*), while atomic force microscopy has enabled the conformation mapping of very short RNA species *in vitro* (*7*). However, the direct visualization of the three-dimensional ultrastructure of mRNA inside cells—where they engage with cellular factors, undergo processing, and transport between subcellular locations—has remained a major challenge in RNA biology.

Electron microscopy has provided insights into the organization of mRNPs (*8*) and of cytoplasmic RNA through polysome imaging (*9*), leading to initial hypotheses of mRNA structure. Fluorescence *in situ* hybridization applied to RNA (RNA-FISH (*10*) and single molecule smRNA-FISH (*11*)) enables transcript-specific visualization of RNA molecules in cells. However, insights into mRNA ultrastructure have been limited to diffraction-limited imaging approaches and to the analysis of one-dimensional segment-to-segment distances within transcripts (*12*, *13*).

Super-resolution microscopy (SRM) (*14*, *15*) has unlocked optical access to the sub-hundred nanometer scale in cell biology, and spurred efforts to determine the distribution and colocalization of proteins in cells and tissue (*16*). Among SRM techniques based on single-molecule localization (SMLM) (*17–19*), DNA-PAINT (*20*) is widely adopted for its multiplexing capabilities (*21–23*), quantitative molecular counting (*24*), and sub-nanometer precision (*25*).

SRM has enabled optical chromatin tracing (*26*), unveiling biological insights of chromatin organization (*27–30*). In contrast, super-resolving an RNA molecule has lagged behind, owing to high RNA-FISH primary probe densities, and stringent resolution requirements. RNA imaging has been carried out using stimulated emission depletion microscopy (*31*), expansion microscopy (*32*, *33*) and DNA-PAINT (*29*, *34*), with recent efforts targeting small RNAs (*35*) through specialized labelling design utilizing nucleotide analogs. While these approaches improved measurements of the cellular distribution of RNA, none have thus far addressed the ultrastructure of single RNAs at sub-transcript resolution.

In this work, we tackle this gap by developing a novel DNA-PAINT combinatorial multiplexing technique, Combi-PAINT, that enables the fast, multiplexed imaging with reduced sample perturbations. We apply Combi-PAINT to mRNA ultrastructure imaging with widefield and MINFLUX SRM modalities, showing the sub-transcript spatial organization of multiple mRNAs in the human cell nucleus and cytoplasm. We find differences in size, shape, and compaction for transcripts of different genetic architecture, which are enabled by a sub-diffraction view of mRNA. Taken together, Combi-PAINT and optical RNA ultrastructure analysis provide the first, to our knowledge, ultrastructural view of mRNA in cells and a generalizable platform for the ultrastructural imaging of any nucleic acid.

## Results

### Combinatorial multiplexing for DNA-PAINT super resolution imaging

DNA-PAINT exploits the transient binding of fluorescent DNA strands (‘imagers’) to their complement (‘handles’), attached to molecules of interest, to achieve the emitter sparsity, or ‘blinking’, necessary for SMLM. However, conventional DNA-PAINT multiplexing requires a number of imaging rounds that scales linearly with the number of desired imaging targets (*21–23*).

To simplify and accelerate high-target number DNA-PAINT imaging, we developed a combinatorial DNA-PAINT multiplexing approach (Combi-PAINT) in which a handle strand, coupled to a target, can recruit more than one imager strand (Handle AB in Fig. 1A). This strategy simplifies DNA-PAINT workflows and increases the number of resolvable targets without increasing the number of imaging rounds (*21*, *36*). At the single-molecule level, the colocalization of signals from multiple imagers encodes for target identity (Fig. 1B). For example, two imagers (A and B) suffice to discriminate three targets (A, AB, B in Fig. 1B, top); by direct generalization to N imagers, a binary mapping enables 2^N^-1 targets. Moreover, allowing arbitrary stoichiometries of imager binding sites on the same handle strand (Fig. 1B, bottom) relaxes the binary mapping constraint, enabling the number of distinguishable handles to grow beyond exponential scaling.

**Fig. 1.**
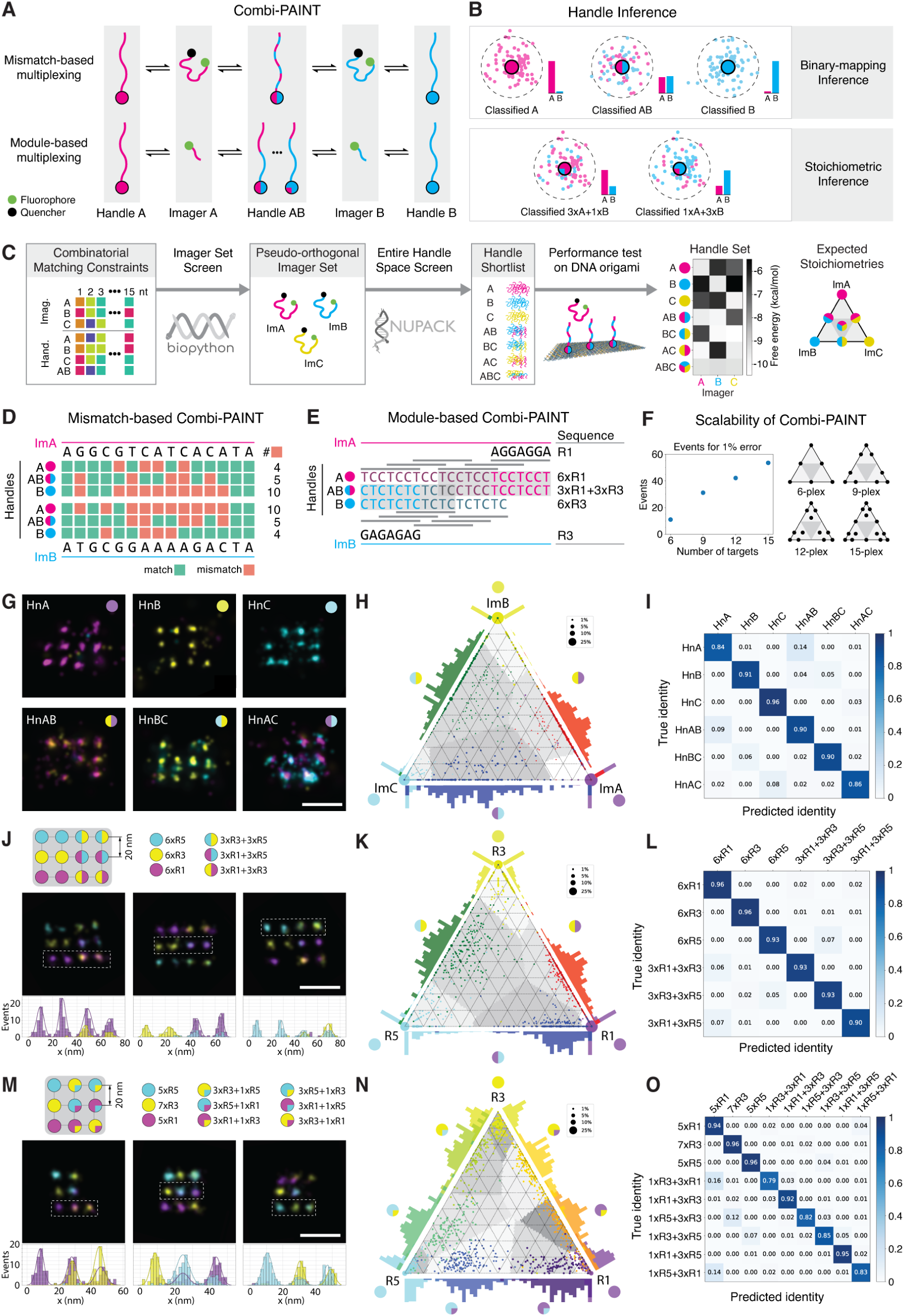
Combinatorial DNA-PAINT multiplexing. (**A**) Concept: *de novo* target DNA handles are designed to bind transiently to several DNA imagers, e.g. handle “AB” binds imagers “A” and “B”. Handle-imager pairs are designed by either (top) distributing base mismatches or (bottom) concatenating short modules. (**B**) The handle inference through classification of clustered localizations can assume either (top) binary-mapping with identical stoichiometries or (bottom) arbitrary stoichiometries of imager mixtures. (**C**) Computational pipeline for designing fluorogenic mismatch-based Combi-PAINT strand families (see Methods). The heatmap on the right shows the binding energy of each imager-handle pair, while the ternary plot shows the position of the handles, represented in the same color scheme as in the matrix, given the proportion of binding to each imager strand. (**D**) Mismatch arrangement of HnA, HnB and HnAB with ImA and ImB. The different arrangement of mismatches between ImA and ImB for HnAB is shown, as well as the much higher number of mismatches for HnA-ImB and HnB-ImA. (**E**) Imager binding pattern of Module-based Combi-PAINT, showing the partial overlap of the binding motifs to reduce handle length. (**F**) Example of scalability of Module-based Combi-PAINT exploiting binding stoichiometry. The plot on the left is a subset of, showing only the optimal scheme for each number of targets. The y-axis represents the number of events needed to achieve a 1% error probability for each target classification in the scheme. The diagrams on the right show the position, in a ternary plot, of each handle species in each scheme. (**G**) 6-color mismatch-based Combi-PAINT; a mixture of DNA origamis with a 20 nm-spaced grid for each handle type were imaged simultaneously over three rounds of exchange-PAINT with each imager: ImA (magenta), ImB (yellow), ImC (cyan). Scalebar: 50 nm. (**H**) Ternary plot representation of classification. Dot coordinates (*p*_*ImA*_, *p*_*ImB*_, *p*_*ImC*_): recorded fraction of each imager per binding site. Color represents ground-truth: HnA (magenta), HnB (yellow), HnC (cyan), HnAB (red), HnBC (green), HnAC (blue). Dot size: fraction of entire population. Target classification is done according to the shaded gray areas, which represent maximum-likelihood classification. DNA-origami for each type: n_HnA_ = 356, n_HnB_ = 424, n_HnC_ = 543, n_HnAB_ = 638, n_HnBC_ = 1026, n_HnAC_ = 199. (**I**) Confusion matrix of the classification. Rows may not sum to 1 due to rounding errors. (**J-L**) Equivalent to (G-I) for Module-based Combi-PAINT for six targets; (J, top) diagrams of origami handle arrangement; (J, bottom) line profile from the localizations in the dashed rectangle. N=136 origamis. Scalebar: 50 nm. (**M-O**) Equivalent to (J-L) for Module-based Combi-PAINT for nine targets. N=253 origamis. Scalebar: 50 nm.

We present two avenues to implement the Combi-PAINT concept (Fig. 1A): (i) a “mismatch-based” design, which exploits engineered base-pair mismatches to yield handle-imager pairs of comparable binding kinetics, and (ii) a “module-based” design, in which handles concatenate short motifs that independently recruit distinct imagers.

Mismatch-based Combi-PAINT requires handle sequences to deviate from exact complementarity, reducing binding stability, which is compensated by increasing sequence length. By choosing the position of base-pair mismatches appropriately, distinct imagers can be programmed to bind to the same handle, allowing combinatorial multiplexing. We devised a computational pipeline for the rational design of such DNA strand families (Fig. 1C, Methods). The pipeline is seeded with randomized DNA sequences constrained by an analytic model of nucleotide dependencies that enforces a binary mapping between imagers and handles (Supplementary Fig. S1). This first *in silico* screen yields pseudo-orthogonal imager sets (ImA, ImB, ImC) and their corresponding handles (HnA, HnB, HnC, HnAB, etc.). As the analytic model does not predict realistic DNA-DNA binding, we produced a second *in silico* screen utilizing NUPACK (*37*). Here, the entire handle space (∼70M combinations) was compared against the chosen imager sets, in search of uniform binding-energies, high off-target rejection and low self-interactions (Supplementary Fig. S2, Supplementary Table S1). Candidate handles were experimentally validated on DNA origami (*38*) by measuring their binding kinetics (Supplementary Table S2). From these, we selected an optimized six-handle set (Supplementary Fig. S3, Supplementary Table S3, Supplementary Table S4, Supplementary Table S5), of which a subset is shown (Fig. 1D) to highlight the mismatch distribution.

We generated families of fluorogenic mismatch-based Combi-PAINT imagers spanning different spectral ranges by appending a quencher at the end of the strand opposite to the fluorophore, thereby suppressing background fluorescence from unbound strands (*39*) (Supplementary Fig. S4, Supplementary Fig. S5, Supplementary Table S5). To validate multiplexing performance, we imaged a mixed sample of six distinct 20-nm grid origamis, one per mismatching handle, over three rounds of Exchange-PAINT imaging (Fig. 1G). For each binding site, identity was inferred from a multinomial distribution model of binding events and compared with the ground truth defined by the origami template (Fig. 1H). The resulting confusion matrix demonstrated high (85-95%) sensitivity even at low coverage (>15 binding events per site; Fig. 1I). In parallel, this experiment provided binding kinetics and photophysical parameters of all imager-handle pairs simultaneously (Supplementary Fig. S6, Supplementary Fig. S7, Supplementary Table S6, Supplementary Table S7). MINFLUX imaging of the nuclear pore complex (*40*) utilizing these imagers exhibited clearly localized clusters (Supplementary Fig. S8).

The second avenue, module-based Combi-PAINT, uses conventional DNA-PAINT handles (*41*) in combined concatenations (Fig. 1A). Given its capability of encoding more than 2^N^-1 handles through arbitrary ratios of binding sites for different imager strands, we aimed to identify optimal stoichiometries. We explored this design space with a maximum-likelihood classifier that computes the classification error as a function of stoichiometry, number of targets, and number of binding events (Supplementary Fig. S9, Supplementary Fig. S10). A subset of this analysis (Fig. 1F) shows the efficiency of the optimal strategies to address 6-, 9-, 12-, and 15-target imaging with three imager strands, which can be decoded with tens of binding events. Increasing the number of imagers to six (*42*), allows up to 63 targets (Supplementary Fig. S11). We employed the three established purine-only imagers to leverage the rapid imaging capabilities of single-class nucleobase strands (*42*) after checking their binding modes (Supplementary Fig. S12).

We characterized the performance of the 6-(Fig. 1J) and 9-(Fig. 1M) target strategies with a similar experiment to the one for mismatch-based Combi-PAINT, but with each individual origami harboring all the handle types. We created a Python pipeline to extract the ground truth identity of the handles from their position on the DNA origami, as well as the number of binding events. The resulting ternary plots (Fig. 1K,N) and confusion matrices (Fig. 1L,O) show high (90-95% and 80-95% for the 6- and 9-color strategies, respectively) sensitivity.

Taken together, we present two strategies, mismatch- and module-based, to perform novel combinatorial DNA-PAINT multiplexing, which we call Combi-PAINT. We show that three imagers can combinatorially encode double and triple the number of targets compared to a non-combinatorial approach. Module-based Combi-PAINT, due to the larger number of available speed-optimized imager strands, presents a higher ceiling to the possible number of addressable targets (Supplementary Fig. S11, Discussion). Mismatch-based Combi-PAINT, on the other hand, excels in background-heavy environments and in the absence of optical sectioning due to the fluorogenicity of its imager strands. Furthermore, while we tested both mismatch- and module-based handles for the purpose of RNA labelling, we found that module-based handles introduced undesired nuclear background under RNA-FISH conditions (Supplementary Fig. S13); however, they remain fully effective for non-nuclear targets and samples that do not undergo FISH hybridization.

### Super-resolved RNA-FISH reveals mRNP morphology

We next integrated Combi-PAINT with RNA-FISH to directly visualize mRNAs in human cells. In conventional smFISH, fluorescent oligonucleotide probes hybridize with complementary mRNA sequences, providing transcript-specific signals at diffraction-limited resolution. In our design, each probe carries two single-stranded DNA extensions flanking the mRNA-binding region: one which engages the smFISH probe, serving as an intrinsic positive controls of RNA presence, and one which supports either the mismatch- or module-based Combi-PAINT readout as described above (*43*) (Fig. 2A).

**Figure 2.**
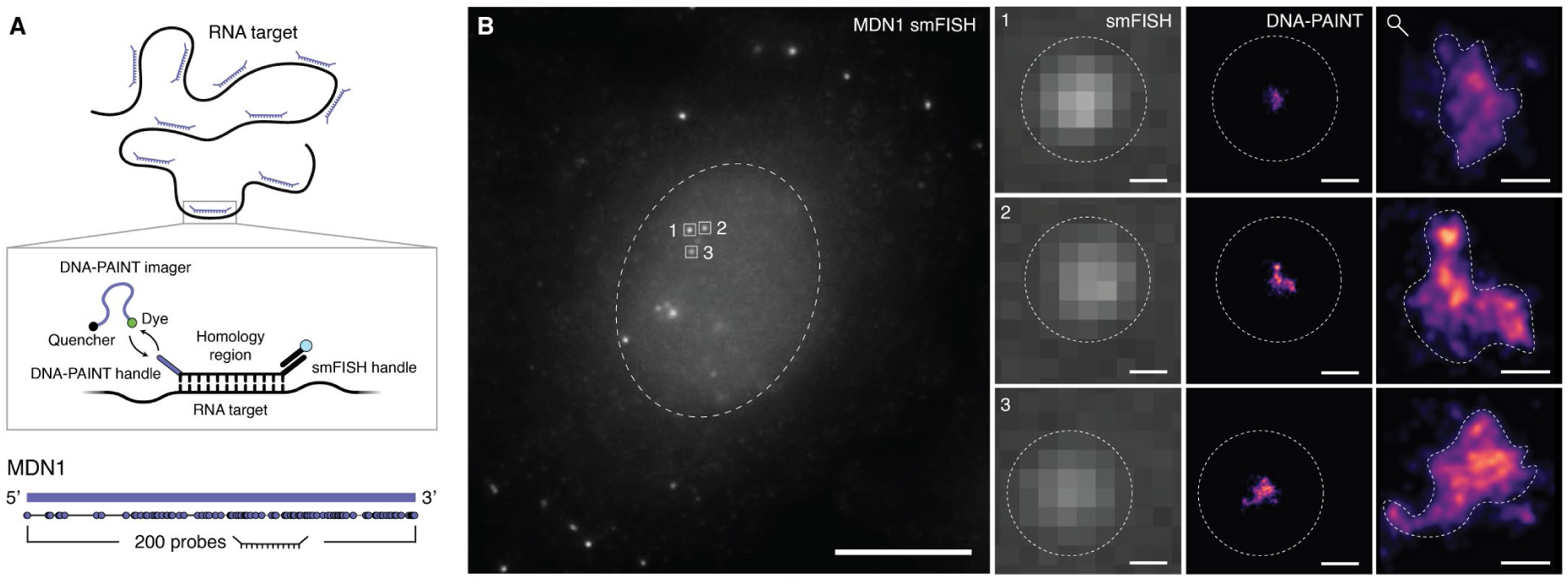
Super resolving mRNA by combining DNA-PAINT and RNA-FISH. (**A**) Cartoon of the RNA-FISH labelling strategy employed and of RNA-FISH probe distribution along the mature MDN1 transcript. (**B**) Left: diffraction-limited single-molecule FISH (smFISH) overview of a cell labelled with RNA-FISH for the MDN1 mRNA. Right, first column: zoom-in of three distinct MDN1 mRNAs marked in the overview; second column: single-color super-resolved DNA-PAINT image at the same scale; third column: zoom-in of DNA-PAINT images with a dashed contour marking the RNP region. Scalebar: 10 μm (cell overview), 200 nm (first and second columns), 50 nm (third column).

As a first demonstration, we targeted MDN1, a well-studied 18 kb-long transcript, which was previously used as a model mRNA due to its large size, moderate copy number in cells and balanced nuclear/cytoplasmic localization to benchmark RNA structure in one dimension (*12*). We designed 200 homology sequences with OligoMiner aiming to sample the mature transcript as uniformly and selectively as possible, and appended with smFISH and mismatch-based Combi-PAINT handles (Methods). The RNA-FISH protocol was optimized for signal-to-background ratio and includes a shielding step (*44*) to reduce nuclear background (Supplementary Fig. S14). These optimizations allowed a clear identification of individual transcripts in high-quality smFISH (Fig. 2B, left) and their super-resolved reconstruction by DNA-PAINT (Fig. 2B, right) with high precision (NeNA (*45*): σ = 4-6 nm), even in the high-background nuclear environment. We excluded rare nuclear multi-transcripts clusters from downstream analysis, which may correspond to transcription sites. Taken together, we directly visualize the morphology of mRNA in two dimensions, for the first time.

To investigate whether different mRNA species follow different ultrastructural rules, we imaged two additional transcripts that differ in their genomic features (Fig. 3A, Supplementary Fig. S15). Alongside MDN1, a long transcript with high intron density (101 splice junctions), we selected AHNAK1, an mRNA of comparable length (18,761 bp) but with low intron density (4 splice junctions, localized at the 5’ end). We also imaged EPRS1, which is of average mRNA length (4,300 nucleotides) but has a high intron density (31 splice junctions). RNA-FISH probes (MDN1: 200, AHNAK1: 98, EPRS1: 46) were uniformly distributed across each mRNA. Each mRNA was then split into three segments containing equal numbers of probes, and the probes of each segment were equipped with the same Combi-PAINT handle. Notably, this arrangement differs from simply interleaving RNA-FISH probes with different conventional DNA-PAINT handles along the transcript, as the stochasticity of probe binding cannot ensure true colocalization of signals, which is a unique feature of Combi-PAINT (Supplementary Fig. S16).

**Figure 3.**
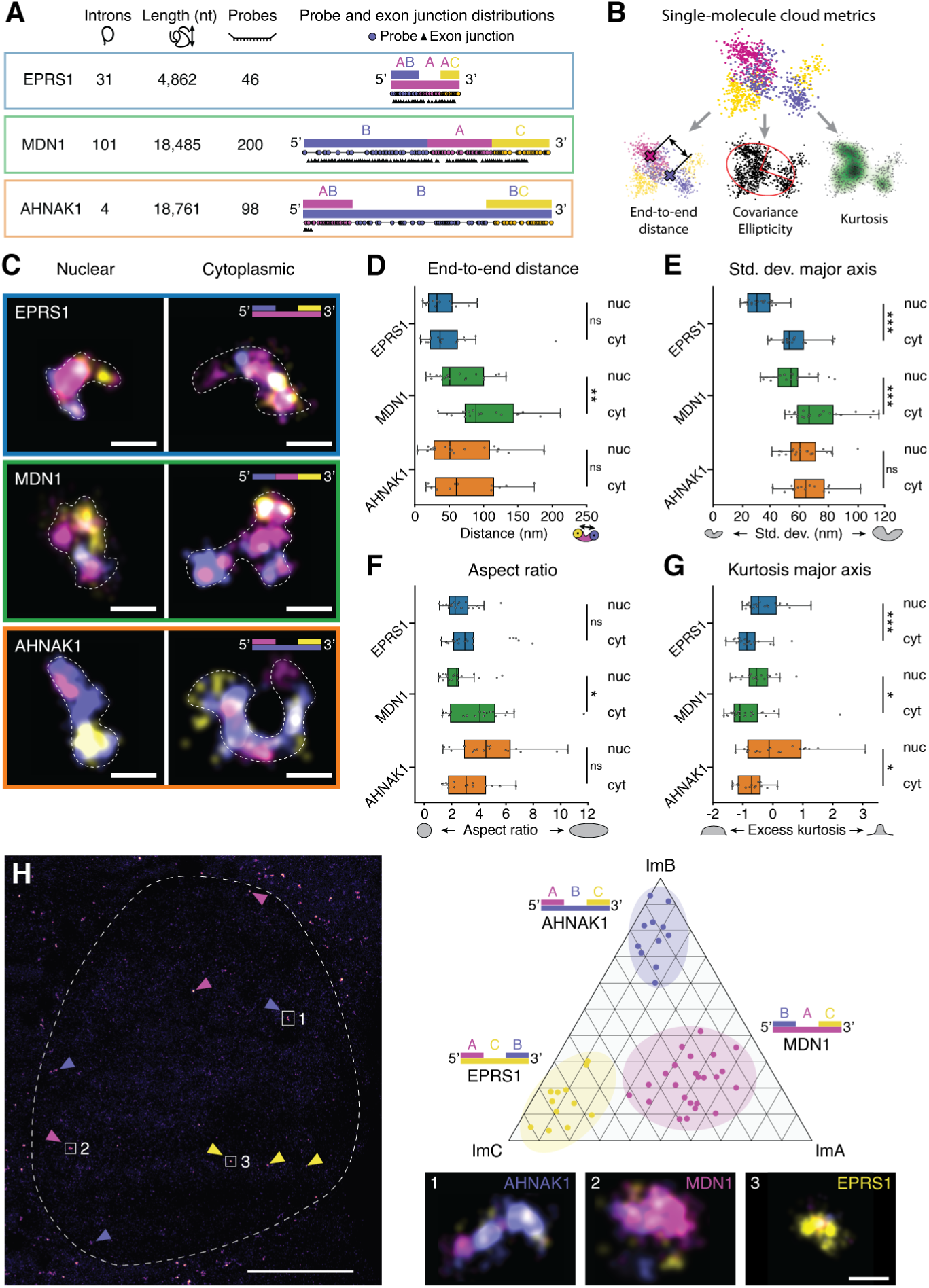
Morphology assessment of distinct mRNAs. (**A**) Overview of the investigated mRNAs’ properties: number of introns, length of the mature transcript, number of designed RNA-FISH probes and sequence map highlighting probe positions (circles) and splice junctions (triangles). (**B**) Extracted metrics from each RNP’s localization cloud. (**C**) Representative combinatorial DNA-PAINT images of RNPs of each of the three genes in the nucleus and cytoplasm, with a dashed contour marking the RNP region. Magenta: ImA, lilac: ImB, yellow: ImC. Scalebar: 100 nm. (**D-G**) Plots of 5’-3’ end-to-end distance (D), major axis standard deviation (E), aspect ratio (F) and major axis kurtosis (G) of the three genes in the nucleus and cytoplasm. *: p<0.05, **: p<0.01, ***: p<0.001, n.s.: non-significant. Data from N>3 experiments per RNA, no. of particles MDN1 nuclear: N=21, MDN1 cytoplasm: N=23, AHNAK1 nuclear: N=20, AHNAK1 cytoplasm: N=17, EPRS1 nuclear: N=22, EPRS1 cytoplasm: N=21. (**H**) Combi-PAINT RNA-FISH image. Left: overview of the cell with nuclear area outlined by the dashed line. A zoom-in of the nuclear numbered particles is shown on the right. Arrowheads show other particles classified in the experiment (magenta: MDN1, lilac: AHNAK1, yellow: EPRS1). Right: ternary plot showing the proportion of binding of each imager strand to each RNP, represented by a single dot. Classification is performed via DBSCAN clustering. Data from N=3 experiments. Scalebars: 5 μm (left), 100 nm (right).

We acquired widefield RNA-FISH DNA-PAINT data over three rounds of Exchange-PAINT separately for each gene (Fig. 3A). The resulting localization clouds were analyzed with a custom pipeline that extracted particle features from single mRNPs (Fig. 3B; Methods). To capture their shapes, we quantified 5’-3’ end-to-end distance, as in classical smFISH, and additional descriptors including covariance-based size and aspect ratio, as well as higher moments such as kurtosis (Fig. 3B). Particles were separated into nuclear and cytoplasmic fractions (Fig. 3C), anticipating morphological changes in the cytoplasm owing to export and translation.

End-to-end distances, which are measurable even without super-resolution, reveal mRNA species-specific behaviors (Fig. 3D). MDN1 showed a pronounced increase in extension from nucleus to cytoplasm, consistent with previous reports (*12*). In contrast, EPRS1 and AHNAK1 exhibited little or no detectable change. Based on this metric alone, one might conclude that MDN1 undergoes ultrastructural changes in the cytoplasm, while the other transcripts remain unchanged.

Super-resolved data provided a richer picture (Fig. 3E-G). Both EPRS1 and MDN1 transcripts increased in major axis size from nucleus to cytoplasm, while AHNAK1 remained unchanged (Fig. 3E). This indicates that transcript length by itself may not dictate the degree of compaction. The aspect ratio analysis revealed another pattern (Fig. 3F): in the nucleus, EPRS1 and MDN1 formed extended globule-like structures, whereas AHNAK1 showed a higher degree of elongation. This indicates that intron density may impact mRNP ultrastructure, consistent with biochemical data indicating a role of the splicing-dependent exon junction complex in mRNP packaging (*4*). The aspect ratios of each mRNP were markedly higher than 1, suggesting that mRNPs adopt different degrees of extended, rod-like structures, consistent with a sequencing-based assay (*46*). A strong change in aspect ratio between nucleus and cytoplasm was observed only for MDN1, while the other transcripts showed similar and unchanged aspect ratios. The lack of change in aspect ratio for cytoplasmic EPRS1 may be attributed to its smaller size, which limits ribosome occupancy and preserved its extended globule-like shape. In contrast, we speculate that the low number of introns of AHNAK1 compared to MDN1 may minimize AHNAK1 compaction in the nucleus. We expect that the imaging of additional RNA species may reveal general rules of mRNA ultrastructure regulation, for example, the relationship between intron density and aspect ratio.

Higher-order moment analysis reinforced these trends (Fig. 3G). Kurtosis distributions showed a consistent shift across the three transcripts: cytoplasmic localization patterns became more tail-heavy, reflecting increased RNA density at the particle periphery. This shift is consistent with prior reports that mRNPs undergo conformational changes in the cytoplasm (*12*). While AHNAK1 did not exhibit detectable changes in the other metrics described above when observed in the nucleus or cytoplasm, it did show a significant change in kurtosis distribution similar to the other transcripts, suggesting that this metric reflects some common event across all transcripts, such as translation.

Finally, we leveraged the combinatorial nature of Combi-PAINT to perform simultaneous RNA-FISH of all three segmented transcripts. Each one was labelled with a different assignment of Combi-PAINT handles, such that one imager binds to the entire transcript, while the other two bind to the 3’ and 5’ ends (see cartoon of labeling scheme, Fig. 3H, right). Over three rounds of imaging (Fig. 3H, left), the different stoichiometry signatures enabled blind assignment of both transcript identity and subregion, as confirmed by ternary plots of binding-event distributions (Fig. 3H, right, Supplementary Fig. S17), showcasing another example of 9-target multiplexing.

Taken together, these results reveal that different human mRNAs adopt distinct ultrastructural properties, some of which are influenced by subcellular localization and processing events.

### Three-dimensional MINFLUX imaging of mRNAs

We next aimed to adapt Combi-PAINT to enable three-dimensional (3D) interrogation of mRNP ultrastructure. Since mRNPs are small, 50 nm to 200 nm particles without a preferential orientation in space, they require an imaging method capable of delivering isotropic, few-nanometer precision in all three dimensions on a densely labeled object—an ability unique to MINFLUX, a single-molecule localization strategy that achieves nanometer precisions in 3D by sequentially interrogating the position of fluorescent emitters with a structured beam of focused light.

The nuclear environment poses challenges for this method due to its high background. (Methods, MINFLUX RNA-FISH imaging). To overcome this, we first optimized the MINFLUX targeted-coordinate-pattern (TCP) size *L* (Fig. 4A) to retrieve high-photon counts, with a large *L* = 150 nm yielding superior localization precision and greater robustness to imperfect convergence onto the fluorophore. Second, axial convergence was achieved through a feedback loop, which proved more resilient to variable out-of-focus background binding events than conventional feedforward approaches. Finally, our custom MINFLUX microscope (*47*) allowed mixing distinct excitation beam shapes within a single MINFLUX cycle; combining donut and top-hat beams for lateral and axial localizations, respectively, provided a more photo-efficient MINFLUX scheme, useful for operation in the noisy nuclear environment (Fig. 4A).

**Figure 4:**
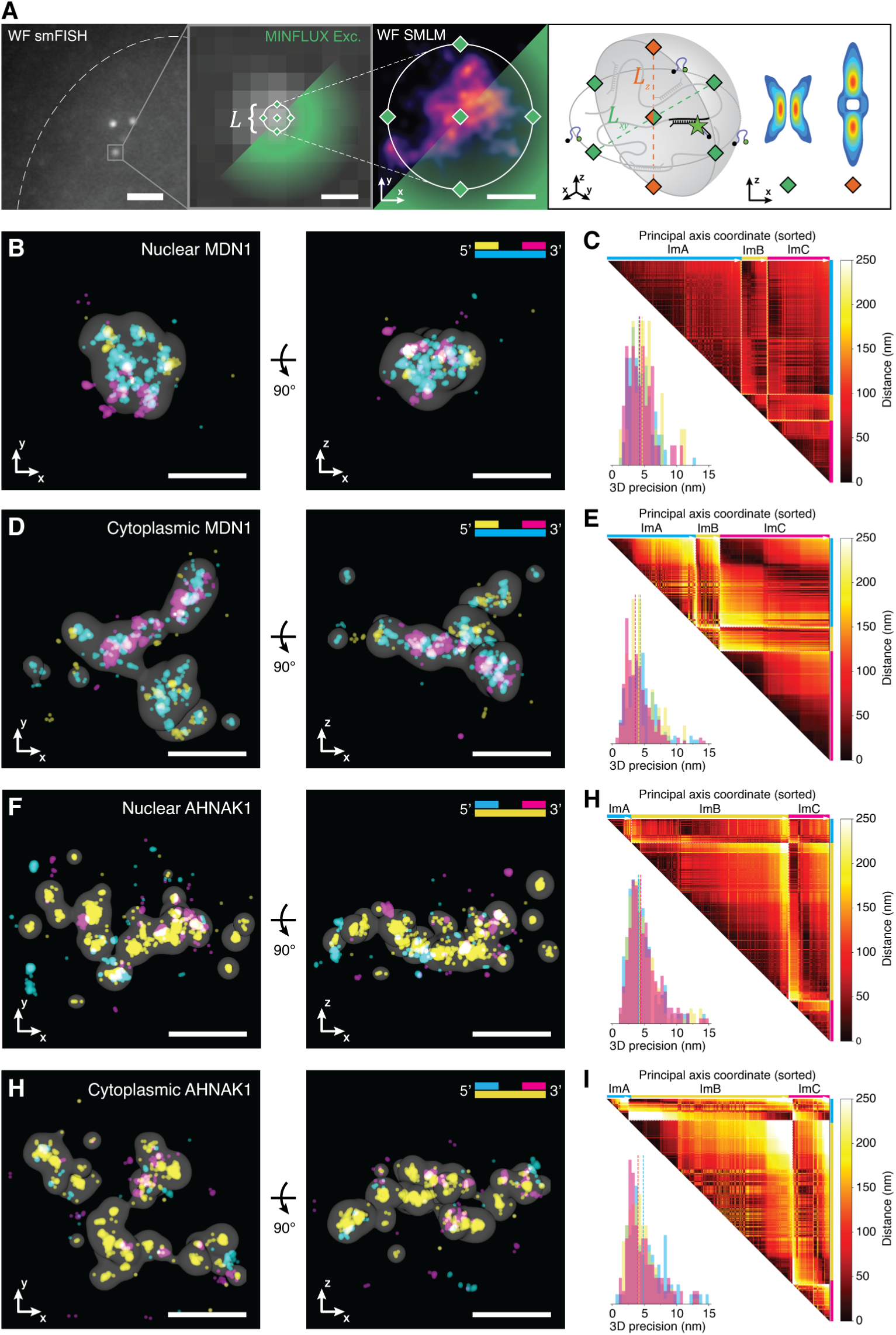
3D MINFLUX imaging of DNA-PAINT RNA-FISH in the nucleus and cytoplasm. (**A**) 3D MINFLUX imaging context. A region of interest chosen by its smFISH signal, is then targeted with iterative 3D MINFLUX imaging. The right panel shows the arrangement of the MINFLUX beams. Green diamond: donut beam, orange diamond: top-hat beam. Scalebars: 2 μm (left), 200 nm (center), 50 nm (right). (**B,D,F,H**) xy and xz projections of single MDN1 RNPs in the nucleus (B) and cytoplasm (D) and of single AHNAK1 RNPs in the nucleus (F) and cytoplasm (H). The shaded volume is a thresholded large-sigma smoothing of the ImA (B,D) or ImB (F,H) rounds, which encompassed the whole MDN1 (B,D) or AHNAK1 (F,H) particles, respectively. Scalebar: 100 nm. (**C**,**E,H,I**) (top) Pairwise distance matrices of the measurements in (B,D,F,H). Each line/column represents a single binding event. Binding events from the same imaging round are grouped together and separated by a dashed line, and each group is internally sorted by first principal component coordinate. (bottom) 3D localization precision achieved in each round of imaging (typically 4-5 nm). Imagers: ImA (cyan), ImB (yellow), ImC (magenta).

With these improvements, we imaged nuclear and cytoplasmic MDN1 and AHNAK1 transcripts *in situ* (Fig. 4B-E, Movies S1-4, Supplementary Fig. S18) at 2 nm localization precision per-axis, corresponding to 4–5 nm resolution in 3D. MDN1 reconstructions revealed a modular arrangement: transcript segments occupy discrete, minimally overlapping volumes, and cytoplasmic particles are overall larger than their nuclear counterparts. This is reflected in the proximity matrices (Fig. 4C,E), which show limited co-occurrence between the ImB and ImC rounds, which labelled the 5’ and 3’ ends respectively and are not expected to colocalize at the single-probe level due to the Combi-PAINT handles distribution. Colocalization of the ImA round with the ImB/ImC rounds is instead expected and can be visually confirmed. In line with widefield MDN1 data the overall size of the particle increases in the cytoplasm. The AHNAK1 reconstructions also confirm the widefield data, showing similar particle sizes and elongation in the nucleus and cytoplasm (Fig. 4F-I). Interestingly, the increased spatial resolution from 3D MINFLUX allows to appreciate a common feature of AHNAK1 cytoplasmic particles, which assume highly curved shapes, with gaps devoid of RNA signal, possibly owing to competition with the translation machinery or other cytoplasmic RNA-binding proteins.

In summary, we introduce a new DNA-PAINT multiplexing method, Combi-PAINT, that is compatible with smRNA-FISH and MINFLUX imaging, allowing us to produce the first low-nanometer-resolution 3D images of nuclear and cytoplasmic mRNA in human cells.

## Discussion

We developed a complete workflow that both advances super-resolution methodology and delivers nanometer-resolved RNA imaging *in situ*. Combi-PAINT enables dense, faster multiplexed labeling, and—paired with iterative 3D MINFLUX—yields segmented reconstructions of human mRNAs and, to our knowledge, the first nanoscopic 3D views of mRNP ultrastructure inside cells. Together, these advances turn studies of mRNA from a diffraction-limited blur into quantifiable particles with shape, size and internal organization.

Combi-PAINT is a general combinatorial multiplexing concept for DNA-PAINT. We implemented it in two ways—mismatch-based and module-based—and established a rational design pipeline that produces families of orthogonal, fluorogenic imagers. Practically, this increases the number of addressable targets without a proportional increase in imagers or exchange cycles, reducing washes, background and drift while remaining compatible with sequential Exchange-PAINT and unmixing approaches based on spectra and fluorescence lifetime (*48*, *49*). In the future, larger libraries of orthogonal, speed-optimized or chemically modified fluorogenic strands should enable scalability to the hundreds, and potentially thousands, of targets (*50*); integration with RESI (*25*), FLASH-/SUM-PAINT (*22*, *23*) and related strategies can further expand resolution and multiplexing; and incorporating additional observables (spectra, lifetime, brightness (*51*)) plus colocalization-aware clustering (e.g., BaGoL inference (*52*)) will harden classification. Because the principle is sequence-agnostic, we expect that Combi-PAINT will generalize to nanoscale chromatin tracing and mapping protein complex stoichiometry and organization.

Super-resolved RNA-FISH coupled to Combi-PAINT reveals transcript-specific particle morphologies in both nucleus and cytoplasm. The level of compaction of mRNAs in the nucleus may correlate with intron density, consistent with a model for exon-junction-complex–mediated packaging of mRNAs (*4*), whereas cytoplasmic particles exhibit a common morphology, likely imposed by translation. Future imaging of many more, and distinct, RNA species will be necessary to investigate global regulatory rules of mRNA ultrastructure. Beyond the traditional end-to-end distance metric, shape descriptors (e.g., covariance-based size, aspect ratio, higher moments such as kurtosis) provide an expanded vocabulary for mRNP ultrastructure, and enable studying the effect of genic features and mRNP composition on the functional mRNA ultrastructure in cells. Notably, the application of DNA-PAINT to RNA we present here is applicable to any RNA target that can be probed by FISH, given enough probes can be generated.

The powerful new toolkit developed here enables the study of RNAs at the nanoscale within intact cells. These methods will facilitate systematic, multi-gene surveys to connect mRNA ultrastructure with gene regulation, cellular function and disease, and to uncover the mechanisms and rules that govern the mRNA life cycle. Similar to how DNA tracing revolutionized our understanding of genome architecture (*27*, *53*), we establish a path to uncover the ultrastructural cell biology of RNA.

## Funding

AP is the recipient of a DOC Fellowship of the Austrian Academy of Sciences at the Institute of Molecular Pathology. PW is the recipient of a Boehringer Ingelheim Fonds (BIF) PhD fellowship. Research in the laboratory of CP is supported by Boehringer-Ingelheim, the European Research Council under the Horizon 2020 research and innovation programme (ERC-2020-STG 949081 RNApaxport) and by the Austrian Science Fund (FWF) doc.funds program DOC177-B (RNA@core: Molecular mechanisms in RNA biology). Research in the laboratory of FB was supported by Boehringer Ingelheim and the European Research Council under the European Union’s Horizon 2020 research and innovation program (ERC-2019-STG 853348 NANO4Life). For the purpose of Open Access, the authors have applied a CC BY public copyright license to any Author Accepted Manuscript (AAM) version arising from this submission.

## Author Contributions

AP and FB conceived the combinatorial multiplexing strategies. CP and FB conceived and identified the application of the super-resolved combinatorial multiplexing strategies for mRNA ultrastructure visualization. AP designed mismatching imaging/handle pairs, designed and folded DNA origami, optimized protocols, performed all DNA origami experiments and their data analysis. AP designed and built the custom widefield optical system. AG and FB built the custom MINFLUX system and programmed the hardware control for the widefield and MINFLUX systems. AP, AG, and FB programmed the MINFLUX processing and visualization software. PW designed RNA-FISH probes, optimized the FISH protocols, performed widefield RNA-FISH experiments and their processing. AP and PW performed MINFLUX RNA-FISH experiments and processing. AP, PW, CP and FB wrote the manuscript. FB and CP supervised the project. All authors proofread and commented on the manuscript.

## Acknowledgements

We thank the Vienna Biocenter Core Facilities (VBCF) for their continuous support, in particular Mathias Madalinski for HPLC purifications, all members of the IMP/IMBA/GMI Mechanical Engineering Center and all members of the IMP/IMBA/GMI BioOptics Facility. We thank the entirety of the Balzarotti and Plaschka labs for discussion and feedback, with special regard to Karina Ayala and Alexander Philipps for help with cell lines. We thank Rupert Faraway and Daniel Zenklusen for helpful discussion regarding the choice of RNA targets. We thank Neos Cruz for sharing and discussing RNA-FISH protocols. We thank April Pawluk (Life Science Editors) and Daniel Zenklusen for critical reading of the manuscript.

## Competing Interests

FB holds patents on principles, embodiments and procedures of MINFLUX.

## Data and Code availability

The raw data supporting the findings of this study are available from the corresponding author upon request. The code for initial imager/handle generation, as well as for exhaustive search and analysis is provided on GitHub at https://github.com/Balzarotti-Lab/fluorogenic_CombiPAINT_handle_generation.

Code for MINFLUX microscope control, MINFLUX data processing and visualization, Combi-PAINT simulations and Combi-PAINT DNA origami processing can be provided upon request.

## Methods

### Widefield microscopy setup

The widefield microscope used for DNA origami experiments was custom built. The light from a Cobolt Jive 150 (0561-04-01-0150-699, Cobolt) was coupled into a polarization-maintaining single-mode fiber (P3-405BPM-FC-5, Thorlabs) using a fiber collimator (60FC-4-M6.2-33, Schäfter+Kirchoff). The output of the fiber was collimated using a 150 mm lens (AC254-150-A-ML, Thorlabs). A cleanup filter (FF01-561/14-25, Semrock) was used to reject any fiber-induced fluorescence. The collimated light was focused by a 400 mm lens (AC254-400-A-ML, Thorlabs) to the back focal plane of a high-numerical aperture oil objective (UPLXAPO60XO, Olympus) after being reflected by a TIRF-suitable dichroic mirror (ZT488/561rpc, Chroma) and passed through a quarter-wave plate (AQWP10M-580, Thorlabs) to induce circular polarization. Between the laser source and the 400 mm focusing lens, a mirror was placed one focal length away from the lens and mounted on a tip/tilt kinematic mirror mount (KM100, Thorlabs), allowing to move from epi-fluorescent illumination to TIRF illumination without displacing the illumination zone. The fluorescence light was passed through a 200 mm 1:1 telescope (2x AC254-100-A-ML, Thorlabs), a 800 nm short-pass filter (FESH0800, Thorlabs) to reject infrared light for the active stabilization, and a dual-notch filter (ZET488/561m, Chroma) to reject residual laser light. The image was formed by focusing with a 150 mm lens (AC254-150-A-ML, Thorlabs) on a PCO panda 4.2 bi camera (Excelitas) or with a 60 mm lens (AC254-060-A-ML, Thorlabs) on PCO edge 2.6 camera (Excelitas) depending on the experiment, leading to a pixel size of 125 and 121 nm respectively. The sample was mounted on a coarse xy stage (M-545.2ML, with M229.25S linear actuators and C-663.11 controllers, Physik Intrumente), itself mounted on a fine xyz stage (P-545.3D8S with E-727 controller, Physik Instrumente), used for active stabilization of the sample. The objective was mounted on a coarse z actuator (L-505.013212F with C-863.12 controller, Physik Instrumente), used for coarse z adjustments. The active stabilization module consists in a darkfield infrared scattering microscope and focus-lock system and is a copy of the one deployed on the MINFLUX setup and described in (*47*). The only modifications in this setup are the z stabilization laser (LuxX 945-200, Omicron) and the coupling/collimation of the z stabilization laser before/after the fiber (60FC-4-A6.2S-02 for both, Schäfter+Kirchoff). The module allows fine (sub-nanometer) and fast (10 Hz) adjustments in xyz of the sample, minimizing short-term drift and maintaining the focus. Residual long term drift (< 25 nm over 1 hour) was corrected in post-processing.

For initial characterization of the imaging strands, as well as widefield DNA-PAINT RNA-FISH, an Elyra 7 microscope (Zeiss) was used. The microscope is equipped with a PCO edge 4.2 camera (Excelitas) and a 63x/1.4 NA plan-apochromat oil immersion objective (Zeiss).

### MINFLUX microscopy setup

The MINFLUX microscope used in this study is described in (*47*).

### Reagents

M13mp18-derived p7249 scaffold (N4040S) was obtained from NEB. Unmodified and biotinylated oligonucleotides (DNA origami staples) were ordered from MWG Eurofins. Azide-modified oligos for conjugation were obtained from Metabion. Other modified oligos for conjugation were ordered from Sigma Aldrich. BHQ2- and BHQ3-modified oligonucleotides were obtained from Biosearch Technologies. IowaFQ-modified oligonucleotides were obtained from IDT. Cy3b-NHS ester (29320) was obtained from Lumiprobe. ATTO488-maleimide (AD 488-41) and ATTO643-maleimide (AD 643-41) were obtained from ATTO-TEC. TCEP hydrochloride (C4706-2G), dimethylformamide (DMF) (227056-100ML), dithiothreitol (DTT) (D0632-5G) and DBCO-PEG4-maleimide (760676-1MG) were obtained from Sigma Aldrich. Absolute ethanol (1009831000) and ammonium chloride (1011451000) were obtained from Merck. Sodium bicarbonate (27778293) was obtained from VWR. Anti-GFP nanobody (N0305) was obtained from NanoTag. Tris-HCl 1 M pH 8, 10x and 1x PBS pH 7.3, 1x TAE pH 8, 1 M HEPES pH 7.3, EDTA 0.5 M pH 8, 1 M magnesium chloride, 1 M potassium chloride and 5 M sodium chloride were obtained from our in-house media kitchen. Pure water was obtained from a MilliQ water filtration system. Neutravidin protein (31000) was obtained from Thermo Scientific. Polyethylene glycol (PEG)-8000 (P2139-500G) and Bovine Serum Albumin-biotin (BSA-biotin) (A8549-10MG) were obtained from Sigma-Aldrich. μ-Slide 8 Well High Glass Bottom (80807) and µ-Slide I Luer Glass Bottom (80177) were obtained from Ibidi. Square #1.5 coverslips (MENZBB024024AC23) and round #1.5 coverslips (6311344) were obtained from Menzel Gläser. Microscope slides (09-3000) were obtained from Bio-Optica. Sail brand single concave glass slides (7103) were obtained from SmartLabs. Hellmanex III (Z805939-1EA) was obtained from Sigma-Aldrich. Double sided tape (665D) was obtained from Scotch. Twinsil speed 22 (1300-1002) was obtained from Picodent. Tween 20 (P9416-50ML), protocatechuate 3,4-dioxygenase from Pseudomonas sp. (PCD) (P8279-25UN), 3,4-dihydroxybenzoic acid (PCA) (506 37580-25G-F), (+−)-6-hydroxy-2,5,7,8-tetramethylchromane-2-carboxylic acid (Trolox) (238813-1G) and sodium hydroxide (30620-1KG-M) were obtained from Sigma-Aldrich. Methanol (c20846.326) was obtained from VWR. Glycerol (A0970-1000) was obtained from BioChemica. 90 nm diameter gold nanoparticles (G-90-100) were obtained from Cytodiagnostics. 40 nm dark-red beads (F8789) were obtained from Invitrogen. Low-glucose DMEM (11880028), FBS (10500064), 100x MEM NEAA (11140035) and trypsin-EDTA (15400054) were obtained from Gibco. Penicillin-streptomycin (P0781) was obtained from Sigma-Aldrich. L-glutamine (25030024) was obtained from Thermo Scientific. 20x SSC (AM9763) was obtained from Invitrogen. Ethylene carbonate (E26258-100G) and Triton X-100 (T8787-100ML) were obtained from Sigma Aldrich. ≥99.5% Formamide (47671) and Dextran sulfate (D8906) were obtained from Sigma Aldrich. 32% Paraformaldehyde (15714) was obtained from Electron Microscopy Sciences. 200 mM VRC (S1402S) was obtained from New England Biolabs.

### Buffers and stock solutions

The buffers used for sample preparation and imaging are the following:

♣ **Buffer A**: 10 mM Tris pH 8.0, 100 mM NaCl. Stored at room temperature.
♣ **Buffer A+**: 10 mM Tris pH 8.0, 100 mM NaCl and 0.05% Tween-20. Stored at room temperature.
♣ **Buffer B+**: 5 mM Tris-HCl pH 8.0, 1 mM EDTA, 12.5 or 50 mM MgCl_2_ and 0.05% Tween-20, optionally supplemented with 1× trolox, 1× PCA and 1× PCD. Stored at room temperature.
♣ **Cellular PAINT buffer**: 1xPBS + 500mM NaCl, supplemented with 1 x PCA, 1 x Trolox and 1 x PCD.
♣ **10x rectangle DNA origami folding buffer**: 50 mM Tris pH 8.0, 10 mM EDTA, 125 mM MgCl_2_. Stored at room temperature.
♣ **10x BSA-biotin**: 10 mg BSA-biotin in 1 ml buffer A. Aliquots stored at −20°C.
♣ **20x Neutravidin**: 10 mg Neutravidin in 1 ml buffer A. Aliquots stored at −20°C.
♣ **40x PCA**: 154 mg PCA in 10 ml water and NaOH, pH adjusted to 9. Aliquots stored at −20°C.
♣ **100x PCD**: 9.3 mg PCD in 13.3 ml of buffer (100 mM Tris-HCl pH 8, 50 mM KCl, 1 mM EDTA, 50% glycerol). Aliquots stored at −20°C.
♣ **100x Trolox**: first, added 100 mg trolox to a mix of 430 μl 100% methanol and 345 μl 1 M NaOH, fully dissolved the trolox, then added 3.2 ml water. pH adjusted to 9. Aliquots stored at −20°C.

### Conjugations

All DNA-PAINT imager strands were labelled in-house using either the maleimide + thiol or the NHS-ester + amine chemistry. For the maleimide + thiol chemistry, 20 nmol of thiol-modified oligonucleotide in 50 μl of 20 mM HEPES pH 7.3 + 100 mM NaCl were added to 50 μl of 2x TCEP (11.5 mg of TCEP hydrochloride in 20 ml of 20 mM HEPES pH 7.3 + 100 mM NaCl, pH adjusted to 7.3 with NaOH) and incubated for 1 hour at room temperature in the dark to reduce the disulfide bonds of the oligonucleotides. Then, 10 μl of 10 mM dye-maleimide in DMF were added to the reaction, the volume was brought to 200 μl with 20 mM HEPES pH 7.3 + 100 mM NaCl, and the reaction was incubated for 2 hours at room temperature in the dark. The reaction was then quenched by adding DTT to a final concentration of 50 mM. For the NHS ester + amine chemistry, 20 nmol of amine-modified oligos were subjected to one round of ethanol precipitation to remove any residual amine from the synthesis process by adding 0.1 volumes of 3 M NaCl and 2.5 volumes of cold absolute ethanol, incubating at - 20°C for 30 min, centrifuging at 21000 g at 4°C for 30 min, and resuspending the pellet in 190 μl of 100 mM sodium bicarbonate pH 8.3. This step was skipped in the case of the 7-nt speed-optimised imagers, as they are too short to be precipitated. Then, 10 μl of 10 mM dye-NHS ester in DMF were added to the reaction, and the reaction was incubated for 2 hours at room temperature in the dark. The reaction was quenched by adding Tris-HCl pH 8 to a final concentration of 50 mM. For both chemistries, the reaction pot after quenching was subjected to one preparative round of ethanol precipitation as just described before purifying via reverse-phase high-performance liquid chromatography on an UltiMate 3000 HPLC system equipped with a Phenomenex Clarity 5 μm oligo-RP column. Again, the preparative ethanol precipitation was skipped in the case of the speed-optimised imagers. The purified oligos were resuspended to 50 μM in 20 mM HEPES pH 7.3 + 100 mM NaCl and stored in low-binding Eppendorf tubes at −80°C. A list of all conjugations performed is provided in Supplementary Table S5.

### DNA origami self-assembly and purification

The two dimensional rectangle origami is based on the design in (*42*). Folding of the DNA origami was accomplished in a 50 μl one-pot reaction consisting of 10 nM scaffold strand, 100 nM unmodified staple strands, excluding the ones substituted by a modified staple strand, 500 nM biotynilated staples and 500 nM modified staple strands (for a list of unmodified, biotynilated and handle-modified staple strands, see Supplementary Material) in 1X folding. The reaction mix was then subjected to a thermal annealing ramp in a thermocycler, consisting in 5 min at 80°C, then cooling immediately to 60°C followed by cooling from 60°C to 4°C in steps of 1°C every 3 min 21 s. The sample was then held at 4°C until purification. In order to remove the excess staple strands and misfolded origamis, the origami was purified via three rounds of PEG precipitation, each consisting in adding the same volume of PEG buffer (1x TAE pH 8.0 + 15% PEG 8000, 500 mM NaCl, 12.5 mM MgCl_2_), centrifuging at 14,000 g at 4°C for 30 min, removing the supernatant and resuspending in 1x folding buffer. The purified origamis were then stored in low-binding Eppendorf tubes at −20°C.

### Bead sample preparation

Glass coverslips were first cleaned via three rounds of water-bath sonication for 10 min in a 2% solution of Hellmanex III. Between each round and at the end, the slides were copiously rinsed with MilliQ water, and finally dried with compressed nitrogen before storage. A flow chamber was formed by placing two double-sided Scotch strips parallel to each other on the microscope slide, and then sandwiching them between the slide and the coverslip. First, 20 μl of a 1:5 dilution of 90 nm gold nanoparticles in 1x PBS was incubated for 5 min in the flow chamber. After a 20 μl wash with 1x PBS, a 1:250 dilution of beads in 1x PBS (previously sonicated for 5 min) was incubated for 10 min. After a 20 μl wash with 1x PBS, the chamber was filled with more 1x PBS, and then sealed with epoxy glue.

### DNA origami sample preparation and imaging

Two kinds of DNA origami samples were prepared, depending on whether the experiment included buffer exchange or not. In case no buffer exchange was needed (fluorogenic handles screening), a flow chamber was made from Scotch double-sided tape sandwiched between the microscope slide and the coverslip, as described previously (*47*). In case exchange was needed (everything else), an Ibidi well slide was used. The following protocol applies to the well slide, with changes for the flow chamber in parentheses. First, the cover glass was cleaned via two 800 μl washes with MilliQ water (Hellmanex cleaning as for bead samples). Then, 200 μl (20 μl) of a 1:4 dilution of 90 nm gold nanoparticles in 1x PBS was incubated for 5 minutes. After a 200 μl (20 μl) wash with 1xPBS and a 200 μl (20 μl) wash with buffer A+, 200 μl (20 μl) of BSA-biotin diluted to 1x in buffer A+ were incubated for 5 minutes. After a 200 μl (20 μl) wash with buffer A+, 200 μl (20 μl) of Neutravidin diluted to 1x in buffer A+ were incubated for 5 minutes. After a 200 μl (20 μl) wash with buffer A+ and a 200 μl (20 μl) wash with buffer B+, the DNA origami solution, diluted to approximately 100 pM in buffer B+, was incubated for 8 minutes. After a 200 μl (20 μl) wash with buffer B+, the imager solution in buffer B+ (imaging buffer) was added to the sample. The imaging buffer was supplemented with 1x PCD, 1x PCA and 1x Trolox. The flow chamber samples were at this point sealed with Twinsil and imaged, while well slides samples were loaded on the microscope, closed with the chamber slide lid, and 15 minutes were waited to allow oxygen scavenging prior to starting to image. Exchange-PAINT was performed by carefully removing the lid to not disturb the actively-stabilized sample, washing with buffer B+ until single-molecule blinking stopped, and finally adding the next imaging buffer before placing back the lid. Details on imaging buffer and imaging conditions are provided in Supplementary Table S2.

### Widefield DNA-PAINT processing

Movies obtained from widefield measurements (DNA origami and cell) were processed with Picasso (*36*). Drift correction of the residual long-term drift was performed via redundant cross-correlation. Alignment of Exchange PAINT rounds was performed via cross-correlation. Linking of the events was performed using a maximum distance of ∼5 times the localisation precision (∼20 nm) and zero dark frames allowed. The processing was also repeated varying these values and without major changes in results. For the mismatch multiplexing experiment, single sites of DNA origami rectangles were manually picked, with the only requirement of the grid being recognizable as of a given species. Since each origami carried twelve binding sites, this requirement was not met only in unfavourable scenarios (non-flatly bound origamis, misfolded origamis, and clusters with several origamis overlapping), making this manual labelling a reliable ground truth classification. If the grid could be recognised, every site present was picked, irrespective of number of events. For modular multiplexing experiments, the whole origami was picked as opposed to single sites. In addition to the previous requirements, the origami was picked only if it did not miss more than three sites, as this would not allow reliable fitting of the template in the subsequent analysis. After picking, custom Python scripts were used to analyze the data. For the mismatch multiplexing experiment, picked sites were filtered from sticking events (> 80% of the localisations happening in a segment <5% of the movie length), selected for repetitive binding (average frame value of the binding events between 10% and 90% of the movie length) and filtered for more than 15 binding events. Notably, all of these procedures are standard for DNA-PAINT, don’t require manual picking (which can be substituted by a clustering algorithm) and can be met by adjusting the imaging time or strand concentration, to meet the required number of events. For the modular multiplexing, origamis were first noise-filtered via a light DBSCAN step with ε ∼ 1.5σ and minimum number of events = 5 to remove isolated events, as well as missing sites. The origamis were then aligned with the template of the designed handle arrangement (as well as the mirror, to account for flipped origamis) using a nearest neighbor distance cost function. The result was visually inspected to remove misfit origamis. This provided the ground truth classification for subsequent analysis. Each binding event was then assigned to a handle via a Gaussian mixture model. Finally, the clusters were selected for more than 10 (6-color multiplexing) or 50 (9-color) total binding events. For both multiplexing strategies, the ground truth classification was used to retrieve the experimental proportion of binding to each imager (the probability vector **p** = (*p*_*ImA*_, *p*_*ImB*_, *p*_*ImC*_)) of each handle species, as well as the average number of binding events for the modular multiplexing. The classification for the mismatch multiplexing experiment was performed, for each site *H*, via choosing the most likely handle species, *H*^∗^, given the formula

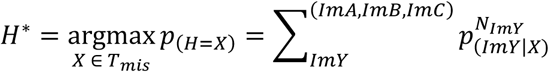

Where *T*_*mis*_ = {HnA, HnB, HnC, HnAB, HnBC, HnAC}, the set of all handles in the 6-color mismatch multiplexing scheme, *p*_(*H*=*X*)_ is the probability of site *H* being handle species X, *p*_(*ImY|*X**)_ the probability of binding of imager *Y* to handle species X, *N*_*ImY*_ the realized number of events from imager *Y*, and the sum being over all three imagers. For the modular multiplexing experiments, a different formula was used to harvest the information contained in the absolute number of binding events, assuming them to be Poisson distributed

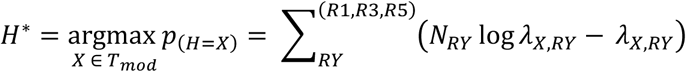

*T*_*mod*_ = {6xR1, 6xR3, 6xR5, 3xR1 + 3xR3, 3xR3 + 3xR5, 3xR1 + 3xR5} for the 6-color modular multiplexing experiment and *T*_*mod*_ = {5xR1, 7xR3, 5xR5, 1xR3 + 3xR1, 1xR1 + 3xR3,1xR5 + 3xR3,1xR3 + 3xR5,1xR1 + 3xR5,1xR5 + 3xR1} for the 9-color modular multiplexing experiment and *λ*_Y_ is the average number of events of imager *Y* for handle species *X*.

### RNA-FISH probe design

FISH-probe libraries were designed for the fully spliced mRNAs of MDN1, AHNAK1 and EPRS1 using OligoMiner (*54*). Homology regions were selected to be between 36 and 41 nt in length with a minimum GC content of 20%. The adjusted melting temperature in 50% formamide and 2xSSC was chosen to fall between 42 and 47 °C. Probes including 5 or more consecutive repeats of any given nucleotide were excluded. The generated probes were further filtered by oligoMiner’s Linear Discrimination Analysis 42°C model with a threshold of 0.5, followed by a jellyfish 18bp k-mer filter with a k of 5. Probes were subsequently appended on their 5’ end with their respective fluorogenic PAINT handle, using a 2-base linker and a stable bio-orthogonal smFISH imager handle (*28*) at their 3’ end. Each mRNA probe set was split into three equal parts by the extension with different PAINT handles with the exact number of probes per mRNA and PAINT handle never falling below 15. Designed probe for each transcript can be found in the Supplementary Material.

### Cell culture

HeLa Kyoto cells (female, HK WT: S. Narumiya (Kyoto University, Kyoto, Japan) were grown in 10 cm cell culture dishes (Falcon) in high-glucose DMEM containing 10% FBS, 100 U/ml penicillin-streptomycin and 1 mM L-glutamine. Cells were kept in a humidified incubator at 37°C and 5% CO_2_. Passaging of cells was conducted every 2-3 days by detachment with 0.05% Trypsin-EDTA. U2OS cells were grown in DMEM without phenol red supplemented with 1x MEM NEAA, 10% FBS, 2 mM L-glutamine and 100 U/ml penicillin-streptomycin. Cells were kept in a humidified incubator at 37°C and 5% CO_2_ and passaged every other day by detachment with 0.05% Trypsin-EDTA. Cells were routinely checked for mycoplasma contaminations.

### Nanobody labelling and purification

Labelling of the anti-GFP nanobody with azide-modified DNA-PAINT handles via DBCO-maleimide crosslinker was performed as previously reported (*23*).

### NUP96 imaging

U2OS NUP96-mEGFP (Cell Line Services no. 300174) were grown for two days on round coverslips to 50-70% confluency. Cells were then rinsed twice with 1xPBS, fixed with 2.4% PFA in 1xPBS for 30 min, rinsed thrice with 1xPBS, neutralized for 5 min with 100 mM NH_4_Cl in 1xPBS, permeabilized for 5 min with 0.25% Triton X-100 in 1xPBS, and incubated for 1 hour in blocking buffer (1 mM EDTA, 3% BSA and 0.02% Tween-20 in 1xPBS). Cells were then incubated with 100 nM anti-GFP nanobody (labelled with a DNA-PAINT handle) in blocking buffer overnight at 4°C under slow shaking. Samples were washed for 5 min with 1xPBS to remove unbound nanobodies, incubated for 5 min with 90 nm gold nanoparticles diluted 1:1 in 1xPBS and rinsed thrice with 1xPBS. Cellular PAINT buffer with DNA-PAINT imager strand diluted to working concentration was deposited in the concave spot of a concave microscope slide, and the seeded coverslip was carefully flipped on top. The chamber was then sealed with Twinsil and imaging carried out similarly to (*55*).

### Design of shielding strands for RNA-FISH

Shielding strands designed to block the DNA-PAINT handles during primary probe incubation but wash in subsequent steps were designed similarly to previous work (*44*). Each mismatch-based Combi-PAINT handle (including linker and padding oligonucleotides) was split in two sub-sequences of melting temperatures as close to each other as possible. The sequences (reverse-complementary to the handle sequence) were ordered and added to the primary incubation mixture as described below.

## RNA-FISH

HeLa Kyoto cells were grown on glass bottom Ibidi-Luer µ-Slides overnight to a confluency of 60-70%. Cells were subsequently washed with 1xPBS for 2 min, followed by fixation with 4% PFA in 1xPBS for 10 min, permeabilization with 0.5% Triton in 1xPBS and a 5 min neutralization in 100 mM NH_4_Cl in 1xPBS. A pre-denaturing step was carried out in which cells were treated with pre-heated 50% formamide/2xSSC and incubated at 60°C for 20 min followed by a 3 min incubation at 78°C. During the pre-denaturation step the hybridization buffer (50% formamide, 10% Dextran sulfate, 2 mM VRC in 2xSSC) was prepared. The pooled library of RNA-FISH-probes was added to the hybridization buffer at a concentration of 7 nM per probe. Shielding strands binding to the DNA-PAINT handles present in the used probe set were added at a concentration of 2 µM each and the mixture was incubated at RT for 20 min. Immediately after finishing the pre-denaturation step cells were incubated with the RNA-FISH-probe containing hybridization buffer over night at 42°C. The next day cells were washed four times with pre-heated 2% Tween in 2xSSC at 60°C before being cooled to RT with 2xSSC for 5 min. A secondary hybridization step for the smFISH imagers was carried out in 5% ethylene carbonate + 2% Tween in 2xSSC with a concentration of 20 nM per smFISH imager for 4 min. This was followed by a wash with 10% formamide + 2% Tween in 2xSSC for 1 min and a final transfer to 1xPBS. Samples were imaged the same day or kept at 4°C for up to two days.

### Widefield RNA-FISH imaging

RNA-FISH samples were incubated with 90 nm gold fiducials (diluted 1:5 in 1xPBS) for 5 min and subsequently washed with 1xPBS three times. Samples were transferred to cellular PAINT buffer with the respective DNA-PAINT imager being added at its desired concentration. Acquisitions were carried out on an Elyra 7 microscope. A plane of interest was chosen by assessing the smFISH signal in the 488 nm channel followed by engaging the definite focus functionality to not lose the chosen focus during the acquisition. Movies were then recorded according to the information in Supplementary Table 1. Exchanges of the DNA-PAINT imagers were carried out by first washing with 1xPBS until no more signal could be detected in the respective imaging channel followed by flowing in a total of 500 µl of the new DNA-PAINT imager-containing cellular PAINT buffer. After ensuring the focused plane had not changed another round of imaging was conducted.

### Widefield RNA-FISH DNA-PAINT processing

Movies obtained from widefield measurements (DNA origami and cell) were processed with Picasso (*36*). Drift correction of the residual long-term drift was performed via redundant cross-correlation and gold fiducials. Localization with a x/y precision above 20 nm were filtered. Alignment of Exchange PAINT rounds was performed via cross-correlation and alignment of gold fiducials. Linking of the events was performed using a maximum distance of ∼3 times the localization precision (∼15 nm) and zero dark frames allowed. The data was subsequently clustered using the DBSCAN implementation in Picasso Render with a minimum number of events of ∼8 and a radius of ∼40 nm. To ensure resulting clusters stem from repeated binding events a custom Python script was used to calculate the average frame value of all events per cluster and discard any that did not fall within a 20% range around the middle of the recorded movie. Following this clusters that overlapped with recorded smFISH signal of the imaged plane were picked and used for further analysis. All further analysis of the widefield data was performed using custom Python scripts. Multiple shape descriptors and parameters were calculated for each RNA species. End-to-end distances were calculated from the centroids of imaging rounds corresponding to the 5’ and 3’ end of each RNA. PCA was carried out and the major axis standard deviation, aspect ratio and major axis kurtosis of each particle were extracted. Significance testing was conducted using a pairwise Mann–Whitney U Test. For particles stemming from multiplexed widefield RNA imaging the ratio of events per exchange PAINT round was calculated and particles were plotted on ternary plots. Subsequent clustered using a DBSCAN implementation was finally used to classify the RNA species per recorded particle.

### MINFLUX RNA-FISH imaging

RNA-FISH samples were incubated with 90 nm gold fiducial beads (diluted 1:3 in 1xPBS) for 5 min and subsequently washed with 1xPBS three times, then they were incubated with 40 nm dark red fiducial beads (diluted 1:10^6^ in 1xPBS) for 10 min and subsequently washed with 1xPBS three times. Samples were transferred to cellular PAINT buffer with the respective DNA-PAINT imager being added at its desired concentration. A typical MINFLUX imaging started with widefield 488 illumination to detect and focus the smFISH signal, and picking of RNA particles for imaging. Widefield 640 nm illumination was then used to pick fluorescent beads for post-processing drift correction and imaging round alignment. Gold nanoparticles for infrared active stabilization were also selected at this point. The stage was then moved by a known amount to compensate for the discrepancy between the position of the 488 nm widefield focus and that of the 640 nm confocal detection and the axial active stabilization was engaged. A typical round of imaging lasted 45 minutes, during which the tip-tilt mirror used for scanning the sample alternated imaging between the RNP and the fluorescent beads, cycling through each every time 20 localizations were acquired (beads) or 5 minutes elapsed (RNP). Exchange rounds were carried out similarly to widefield experiments. The maximum laser power was approximately 40 μW at the sample plane.

Mixed-beams iterative MINFLUX imaging was performed similarly to previous works (*47*, *55*). The 3D MINFLUX strategy consisted of:

| Iteration number | Beam arrangement | Relative power | Live estimator $\beta_0, \beta_1$ | $L_{xy}$ (nm) | $L_z$ (nm) | Number of photons | Minimum time (ms) | Maximum time (ms) |
| --- | --- | --- | --- | --- | --- | --- | --- | --- |
| 0 | 7 G | 0.15 | 1, 2 | 400 | - | 200 | - | - |
| 1 | 4 G | 0.15 | 1, 1 | 300 | - | 200 | - | - |
| 2 | 5 V | 0.25 | 0.8, 6 | 150 | - | 300 | - | - |
| 3 | 3 TH | 1 | 0.4, 1.5 | - | 214 | - | 15 | 20 |
| 4 | 5 V + 3 TH | 1 | 0.8, 22 | 150 | 214 | 30000 | - | 400 |

7 G: 6 Gaussians on a hexagon’s vertices, plus one at the center; 4 G: 4 Gaussians on a square’s vertices; 5 V: 4 vortices on a square’s vertices, plus one at the center; 3 TH: 3 top hats on a line along the axial direction.

Iteration 3 was controlled by a proportional-integral feedback loop to make convergence in z more robust, using the live estimated molecule’s position at each MINFLUX cycle as setpoint of the loop and acting on the top hat’s interferometric deflection. This iteration did not stop upon collecting a certain number of photons, but instead stopped upon reaching a maximum allowed time or upon reaching the setpoint (with a tolerance of 5 nm) after a minimum time. The proportional-integral gains of the feedback loop were optimized on DNA origamis harboring a single, permanently-bound ATTO647N molecule (*47*) and additional free-floating ATTO643, at a concentration that simulated the nuclear Signal-to-Background Ratio (SBR), aiming for balanced settling time and overshooting. Iteration 4 was designed in terms of L and photon number to extract a large number of photons from the molecule, as this is optimal for localization precision at low signal-to-background ratios. Furthermore, isotropy of the localization precision in the 3 spatial directions was optimized by using 4 times more exposure time for the z (top hat) exposures as it is for the xy (vortex) exposures.

### MINFLUX processing

The processing of the MINFLUX data acquired in this work was achieved via a Python implementation of previously used algorithms (*47*, *56*). Briefly, the photon trace of all MINFLUX last iterations was segmented into segments of a defined number of photons (typically 7500). These segments were then localised using an MLE estimator. Filtering of the localizations was done based on *p*_*0xy*_(*n*_0,*vortex*_⁄*N*_*vortex*_, the ratio of the photons from the central vortex in the last iteration and all vortex exposures in the last iteration, typically chosen to be < 0.15), *p*_0*z*_(*n*_*0,top hat*_⁄*N*_*top hat*_, the ratio of the photons from the central top hat in the last iteration and all top hat exposures in the last iteration, tipically chosen to be < 0.25) and SBR (tipically chosen to be > 0.5) (Supplementary Fig. S19). Fluorescent bead localizations were then used to perform correction of residual drift and channel alignment: localizations coming from each bead at each timepoint of an imaging round were averaged to generate a high-precision localization. The trace formed by these localizations was then fit with a third-degree spline to interpolate the drift between timepoints. This continuous drift trajectory was then used to correct each MINFLUX localization for drift. The shift between rounds was similarly corrected by calculating the shift between the same bead in subsequent rounds of imaging and subtracting it to the entire round. Visualization of the data was done via Napari (*57*) after binning the 3D data (creating a 3D voxelated image) with a bin (voxel) size of 1.5-2 nm and applying a gaussian blurring of σ equal to the localization precision of the dataset (typically 2-2.5 nm). This localization precision was estimated in a similar way to the NeNA precision widely used in DNA-PAINT works, by retrieving the distribution of distance between consecutive localizations of the same binding event.

### Fluorogenic mismatch-based Combi-PAINT strand design and screening

The design of the fluorogenic mismatch-based Combi-PAINT strands was done entirely in Python, following the steps in Fig. 1B. A mathematical model (depicted in Supplementary Fig. S1) was created to allow the design of imager-handle pairs of the desired length, number of mismatches (15 nt and 4 respectively, in agreement with results from (*39*)) when binding is desired (e.g., ImA to HnA), and as many mismatches as possible when it is not (to minimize crosstalk, e.g., ImA to HnB). The Python program assigned random bases to the nucleotide positions described in Supplementary Fig. S1, and implemented a random shuffling of all bases apart from the first and last, as binding at the ends is desired anyway to make sure dye and quencher are truly far apart. The shuffling is necessary as the specific arrangement of matches and mismatches influences the strength and efficiency of binding, or in other words, the kinetics of binding. The program then runs the following checks using BioPython functions (*58*):

1. Pairwise local alignment of each undesired imager-handle pair (to avoid crosstalk)
2. Pairwise local alignment of each imager-imager pair (to avoid dimers)
3. Pairwise local alignment of each imager strands’ start and end (to avoid hairpins)
4. Melting temperature (to match a desired range)

Pairwise local alignments scored matches as +1, mismatches as 0, and gap initiation and elongation as −1. Necessary scores to pass were <6, <7 and <4 for check 1, 2 and 3 respectively. The accepted melting temperature range at check 4 (calculated using DNA_NN3 nearest neighbor thermodynamics for 5 mM Tris, 50 mM Na^+^ and 10 mM Mg^2+^) was −45°C < < −25°C, but allowing up to 4 of the 12 desired pairwise bindings to lie outside this range. If these checks are passed, the imager-handle set is passed to NUPACK (*37*) to calculate all pairwise binding energies. The choice of the initial set was done by reviewing these energies for weakness (fast binding) and uniformity (similar binding time). The exhaustive search for optimal handles was again done entirely in Python using NUPACK. In this case, the imagers were left fixed, and all 4^13^ oligonucleotide handle sequences were generated (the first and last nucleotide were left fixed and matching). The following checks were ran:

1. Intramolecular binding energy (needs to be weaker than a threshold)
2. Binding energy with each imager (has to be outside certain ranges)

The intramolecular binding energy (calculated for 25°C, 50 mM Na^+^ and 10 mM Mg^2+^ with model “dna04-nupack3” and ensemble “some-nupack3”) was set to be higher than −0.8 kcal/mol. The binding energy range to exclude (calculated in the same conditions as the intramolecular binding) was < −10 kcal/mol and −8.5 < < −7 kcal/mol. The first range was too strong a binding (desired or not), while the second range a binding too weak if desired and too strong if undesired. If a handle passed these checks, depending on which binding energies to the imagers were suitable, the handle was classified as a candidate for its handle type (e.g., if the binding energy to ImA and ImB were outside the excluded ranges, the handle is classified as a candidate HnAB). The handle sequences and binding energies were then filtered by species (the list of filters is provided in Supplementary Table S1 until ∼100 sequences per species were left. For these sequences, the secondary structure of binding to their strands was predicted with NUPACK, and visually inspected for in-frame binding and strong binding at the extremities, with preference for weaker-binding handles. The top 10 handles of each handle species were given a 3’, 2-nucleotide linker and a 5’ 1-nucleotide overhang (*39*), designed to not modify the binding to the imagers or the intramolecular binding, and measured individually on DNA origamis to extract kinetics and evaluate image quality to select the best-performing handle of each species. Remarkably, all HnA, HnB and HnC candidates, and the majority of HnAB, HnBC and HnAC candidates yielded good-quality images, proving the effectiveness of the computational pipeline in finding functional handle sequences. HnABC was excluded for this work, since it would not be possible to distinguish from a cellular feature binding all imagers or the fluorophore indiscriminately. The code for initial imager/handle generation, as well as for exhaustive search and analysis is provided on GitHub.

### Simulation of stoichiometries for modular multiplexing

Simulation of optimal stoichiometries for modular multiplexing was performed numerically in Python. Assuming three imager strand types *A*, *B*, *C* (in our case, R1, R3, R5), we modelled modular Combi-PAINT sets composed of handles each built with 3, 4, 5 or 6 possibly mixed concatenations of such types (3x*A* and 2x*A*+1x*B* could for example be part of the same set, but 5x*A*+1x*B* cannot). Only combinations that are radially symmetric on the ternary triangle were considered, as these better cover the ternary plot’s space. Under these constraints, there are 6 Combi-PAINT handle sets for 3 modules, 12 for 4 modules, 35 for 5 and 272 for 6.

After defining a Combi-PAINT handle set, its performance is evaluated by calculating, for each of its handles, the classification error probability assuming *N* binding events. *N* = *n*_*A*_ + *n*_*B*_ + *n*_*C*_, the sum of binding events from each imager strand, was sampled logarithmically. We also define *D*_*N*_ = {(*n*_*A*_, *n*_*B*_, *n*_*C*_) | *n*_*A*_ + *n*_*B*_ + *n*_*C*_ = *N*} as the set of all possible combinations of binding event numbers that sum to *N*. Then, the error probability for handle species X is defined as

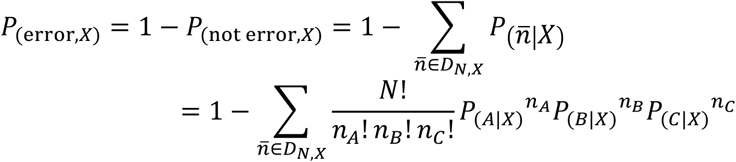

Where *n̄* is a given realization of *n*_*A*_, *n*_*B*_, *n*_*C*_ events from the imagers, *D*_*N*,*X*_ ⊆ *D*_*N*_ is the set of elements *n̄* for which handle species X is the most likely to have originated *n̄*, in analogy to the classification problem in the Widefield DNA-PAINT processing section, and *P*_(*A*|*X*)_, *P*_(*B*|*X*)_, *P*_(*C*|*X*)_ are the probabilities of binding of *A*, *B*, *C* assuming handle species X as the target. For each handle in a combination, the number of events necessary to obtain a reference error probability (1%) was determined by interpolation of the calculated *N* values, and the maximum value across handles of a given combination was retrieved as a metric of performance of the entire handle set.

### Measurement of fluorogenicity factor

The fluorogenicity factor was measured on a LSM980 Axio Observer (Zeiss) inverted confocal laser scanning microscope. Samples of the fluorogenic strand alone (0.5 μM) or with an excess of the reverse complementary oligo (12.5 μM) in buffer B+PCA/PCD/trolox (if the dye was Cy3b, no additives for the far-red dyes) were made in separate flow chambers. The fluorogenic strand solution for each imager was made once and was used for both samples, to avoid pipetting error. The samples were measured at the same focus depth, several microns beyond the coverglass surface. All relevant parameters (confocal pinhole size, detector gain, detection spectral range, pixel dwell time) were kept constant between measurement, while the laser power was adjusted in order to not saturate the detector when measuring the brighter sample with reverse complement, and to fully place the histogram of pixel values above the detector baseline when measuring the dimmer sample without reverse complement. The linearity of the laser power versus % value was ensured prior to the measurement. The average pixel value over the image was then calculated and normalized to the laser intensity used.

