## Supplementary Information for "Super-resolved imaging of mRNA ultrastructure in cells"

**Supplementary Notes**

Supplementary Table S6 Association rate constant between fluorogenic imager-handle pairs. 30
Supplementary Table S7 Dissociation rate constant between fluorogenic imager-handle pairs. 31

### Supplementary Notes

**Supplementary Note 1 Speed considerations of Combi-PAINT**

Combinatorial multiplexing strategies are, in general, utilized to economize a scarce resource like the required measurement time, the number of readouts, the required energy, or the total signal available. In this note, we compare the required imaging time of standard DNA-PAINT multiplexing (non-combinatorial: NC) versus Combi-PAINT (combinatorial: C) under the constraint of equal probability of "double binding" event.

We define:

| Symbol | Definition |
|---|---|
| $H$ | Number of handles or targets |
| $I$ | Number of imagers |
| $h \in \{1, \dots, H\}$ | Index for handles |
| $i \in \{1, \dots, I\}$ | Index for imagers |
| $\mathrm{Hn}_h$ | Handle names |
| $\mathrm{Im}_i$ | Imager names |
| $k_{\mathrm{on},i,h} = k_{\mathrm{on}}$ | Molar binding rate of imager $\mathrm{Im}_i$ to handle $\mathrm{Hn}_h$ |
| $C_i$ | Concentration of imager $\mathrm{Im}_i$ |
| $C_{\mathrm{max},i,h}$ | Maximum tolerable concentration |
| $N_i$ | Number of binding events of imager $\mathrm{Im}_i$ |
| $t_i$ | Imaging time for imager $\mathrm{Im}_i$ |
| $T$ | Total imaging time |

Remarks:

- For the effects of this computations, we assume identical values of $k_{\mathrm{on}}$ for all imagers and handles.
- The non-combinatorial case corresponds to $I = H$, while the combinatorial case has $I < H$.
- We consider that the statistics of DNA binding events follows a Poisson distribution, with the average number of binding events $N$ in a time $t$ with an imager concentration $C$ and a molar binding rate $k_{\mathrm{on}}$ yielding

$$N = k_{\mathrm{on}} C t \tag{1}$$

**Non-combinatorial case**

Here, the number of handles is equal to the number of imagers, $I = H$, therefore, the experiment requires $H$ imaging rounds. From eq. (1), a single imaging round yields $N_i = k_{\mathrm{on}} C_i t_i$ binding events and the total imaging time is

$$T^{(\mathrm{NC})} = \sum_{i=1}^{I} t_i = I t_i = H t_i \tag{2}$$

where equal time is assumed for all imager rounds. This expression can also be written as

$$T^{(\mathrm{NC})} = \sum_{i=1}^{I} \frac{N_i}{C_i k_{\mathrm{on}}} = \frac{\sum_{i=1}^{I} N_i}{C k_{\mathrm{on}}} = \frac{H N_i}{C k_{\mathrm{on}}} \tag{3}$$

where $C$ is the concentration of any imager and $I = H$ was utilized.

**Comparison constrained by the probability of "double binding" event occurrence**

We now focus on the case of high density, where many handle strands fall within the same diffraction-limited area, making them indistinguishable without SRM.

Defining $C_{\max}$ as the imager concentration that yields an acceptable probability of "double binding" events within a diffraction-limited area, the total imaging time of the non-combinatorial case of eq. 3 can be written as

$$T^{(\mathrm{NC})} = \sum_{i=1}^{H} \frac{N_i}{C_{\max,i}^{(\mathrm{NC})} k_{\mathrm{on},i}^{(\mathrm{NC})}} = H t_i^{(\mathrm{NC})} \quad (4)$$

For the combinatorial strategy, we define the density of each target species $h$ as $\delta_h$ and, without loss of generality to make our point, we assume $\delta_h = \delta$. Likewise, we define $d_i = d$, the density of the target handles that bind any given imager, or "imager density".

In analogy to the pigeonhole principle, the combinatorial strategy effectively increases the "imager density" $d$ by using less imaging strands than the non-combinatorial strategy. The specific ratio of imager density between both strategies, $d_{\mathrm{C}}/d_{\mathrm{NC}}$, depends on the specific combinatorial multiplexing scheme (see specific example below) and is equal to $H/I$. The equation for the imaging time per imager, in the combinatorial case, becomes

$$t_i^{(\mathrm{C})} = \frac{N_i^{(\mathrm{C})}}{C_{\max,i}^{(\mathrm{C})} k_{\mathrm{on},i}^{(\mathrm{C})}} = \frac{N_i^{(\mathrm{NC})}}{\left(C_{\max,i}^{(\mathrm{NC})} d_{\mathrm{NC}}/d_{\mathrm{C}}\right) k_{\mathrm{on},i}^{(\mathrm{NC})}} = \frac{d_{\mathrm{C}}}{d_{\mathrm{NC}}} t_i^{(\mathrm{NC})} = \frac{H}{I} t_i^{(\mathrm{NC})} \quad (5)$$

Here, we assume $N_i^{(\mathrm{C})} = N_i^{(\mathrm{NC})}$ and $k_{\mathrm{on},i}^{(\mathrm{C})} = k_{\mathrm{on},i}^{(\mathrm{NC})}$ because they are determined by the desired image quality and imager strand thermodynamics, which are not dependent on the chosen multiplexing strategy. In contrast, $C_{\max,i}^{(\mathrm{C})} = C_{\max,i}^{(\mathrm{NC})} d_{\mathrm{NC}}/d_{\mathrm{C}}$ because the increased imager strand binding site density imposes a proportional reduction in imager strand concentration to prevent overlapping blinking events. This leads to a proportional increase in imaging time. The equation for total imaging time then becomes

$$T^{(\mathrm{C})} = I t_i^{(\mathrm{C})} = I \frac{H}{I} t_i^{(\mathrm{NC})} = H t_i^{(\mathrm{NC})} = T^{(\mathrm{NC})} \quad (6)$$

Therefore, a speed advantage is obtained by having to perform less imager exchange rounds, which are especially time-consuming in dense, large samples like tissues, and goes from 2.5x less wash time (6-color NC imaging, 5 washes, versus 6-color C imaging, 2 washes), 4x less wash time (9-color NC imaging, 8 washes, versus 9-color C imaging, 2 washes) to 12.2x less wash time (62-color NC imaging, 61 washes, versus 62-color C imaging, 5 washes, Supplementary Fig. S11). Furthermore, if information about the spatial arrangement of the targets is available, Combi-PAINT provides a speed advantage uniquely due to multiplexing. For example, if one target is known to have a nuclear localization, while a second target is known to have a cytoplasmic membrane localization, these targets are not expected to lead to overlapping blinking events, even at high imager concentrations. Both targets can then bind to a given imager in a Combi-PAINT multiplexing scenario without the density of binding sites for that imager increasing, leading to a modification of eq. (7) that yields a strict improvement in imaging time.

This is analogous to the decrease in imaging time in multiplexing schemes like MERFISH compared to regular, non-combinatorial FISH: the spatial separation between particles allows the problem to be reduced to a pure decoding problem, in which the advantage of a combinatorial strategy is exactly equal to the reduction in rounds of imaging, $H/I$.

| | **Non-combinatorial label** | **Combinatorial label** |
|---|---|---|
| Target 1 | 2xR1 | 2xR1 |
| Target 2 | 2xR2 | 1xR1+1xR2 |
| Target 3 | 2xR3 | 2xR2 |
| $K$ | 3 | 3 |
| $I$ | 3 | 2 |

#### Supplementary Figures

| | 1 | 2 | 3 | 4 | 5 | 6 | 7 | 8 | 9 | 10 | 11 | 12 | 13 | 14 | 15 |
|---|---|---|---|---|---|---|---|---|---|---|---|---|---|---|---|
| ImA | $X_{1ABC}$ | $X_{2A}$ | $X_{3A}$ | $X_{4A}$ | $X_{5A}$ | $X_{6AC}$ | $X_{7AC}$ | $X_{8AC}$ | $X_{9AC}$ | $X_{10AB}$ | $X_{11AB}$ | $X_{12AB}$ | $X_{13AB}$ | $X_{14ABC}$ | $X_{15ABC}$ |
| ImB | $X_{1ABC}$ | $X_{2BC}$ | $X_{3BC}$ | $X_{4BC}$ | $X_{5BC}$ | $X_{6B}$ | $X_{7B}$ | $X_{8B}$ | $X_{9B}$ | $X_{10AB}$ | $X_{11AB}$ | $X_{12AB}$ | $X_{13AB}$ | $X_{14ABC}$ | $X_{15ABC}$ |
| ImC | $X_{1ABC}$ | $X_{2BC}$ | $X_{3BC}$ | $X_{4BC}$ | $X_{5BC}$ | $X_{6AC}$ | $X_{7AC}$ | $X_{8AC}$ | $X_{9AC}$ | $X_{10C}$ | $X_{11C}$ | $X_{12C}$ | $X_{13C}$ | $X_{14ABC}$ | $X_{15ABC}$ |
| HnA | $Y_{1ABC}$ | $Y_{2A}$ | $Y_{3A}$ | $Y_{4A}$ | $Y_{5A}$ | $Y_{6AC}$ | $Y_{7AC}$ | $Y_{8AC}$ | $Y_{9M}$ | $Y_{10AB}$ | $Y_{11AB}$ | $Y_{12M}$ | $Y_{13M}$ | $Y_{14M}$ | $Y_{15ABC}$ |
| HnB | $Y_{1ABC}$ | $Y_{2BC}$ | $Y_{3BC}$ | $Y_{4M}$ | $Y_{5M}$ | $Y_{6B}$ | $Y_{7B}$ | $Y_{8B}$ | $Y_{9B}$ | $Y_{10AB}$ | $Y_{11AB}$ | $Y_{12AB}$ | $Y_{13M}$ | $Y_{14M}$ | $Y_{15ABC}$ |
| HnC | $Y_{1ABC}$ | $Y_{2BC}$ | $Y_{3BC}$ | $Y_{4BC}$ | $Y_{5M}$ | $Y_{6AC}$ | $Y_{7AC}$ | $Y_{8M}$ | $Y_{9M}$ | $Y_{10C}$ | $Y_{11C}$ | $Y_{12C}$ | $Y_{13C}$ | $Y_{14M}$ | $Y_{15ABC}$ |
| HnAB | $Y_{1ABC}$ | $Y_{2A}$ | $Y_{3A}$ | $Y_{4A}$ | $Y_{5A}$ | $Y_{6B}$ | $Y_{7B}$ | $Y_{8B}$ | $Y_{9B}$ | $Y_{10AB}$ | $Y_{11AB}$ | $Y_{12AB}$ | $Y_{13AB}$ | $Y_{14ABC}$ | $Y_{15ABC}$ |
| HnBC | $Y_{1ABC}$ | $Y_{2BC}$ | $Y_{3BC}$ | $Y_{4BC}$ | $Y_{5BC}$ | $Y_{6B}$ | $Y_{7B}$ | $Y_{8B}$ | $Y_{9B}$ | $Y_{10C}$ | $Y_{11C}$ | $Y_{12C}$ | $Y_{13C}$ | $Y_{14ABC}$ | $Y_{15ABC}$ |
| HnAC | $Y_{1ABC}$ | $Y_{2A}$ | $Y_{3A}$ | $Y_{4A}$ | $Y_{5A}$ | $Y_{6AC}$ | $Y_{7AC}$ | $Y_{8AC}$ | $Y_{9AC}$ | $Y_{10C}$ | $Y_{11C}$ | $Y_{12C}$ | $Y_{13C}$ | $Y_{14ABC}$ | $Y_{15ABC}$ |
| HnABC | $Y_{1ABC}$ | $Y_{2BC}$ | $Y_{3BC}$ | $Y_{4BC}$ | $Y_{5BC}$ | $Y_{6AC}$ | $Y_{7AC}$ | $Y_{8AC}$ | $Y_{9AC}$ | $Y_{10AB}$ | $Y_{11AB}$ | $Y_{12AB}$ | $Y_{13AB}$ | $Y_{14ABC}$ | $Y_{15ABC}$ |

**Supplementary Fig. S1 Combinatorial model for mismatching handle-imager generation.**

Combinatorial model to achieve combinations via mismatches. Each symbol in the table is a DNA nucleotide that obeys the following rules: 1) $X_{i\alpha} = \mathrm{complement}(Y_{i\alpha})$, where complement is the Watson-Crick complement; 2) $X_{i\alpha} \neq X_{i\beta}$; 3) $X_{i\alpha}$ and $X_{j\alpha}$ may or may not be the same nucleotide; 4) $Y_{iM} \neq Y_{i\alpha}$ and $Y_{iM} \neq Y_{i\beta}$, so $Y_{iM}$ is a mismatch with all imagers. The model allows to have 4 mismatches between each binding imager-handle pair, and the maximum possible number of mismatches (>9) between non-binding imager-handle pairs. Handle sequences need to be reversed to obtain the 5'→3' handle sequence. The table columns are ordered to show the symmetries of the model, but should be shuffled to explore different mismatch positions along the sequence space, which influence binding kinetics.

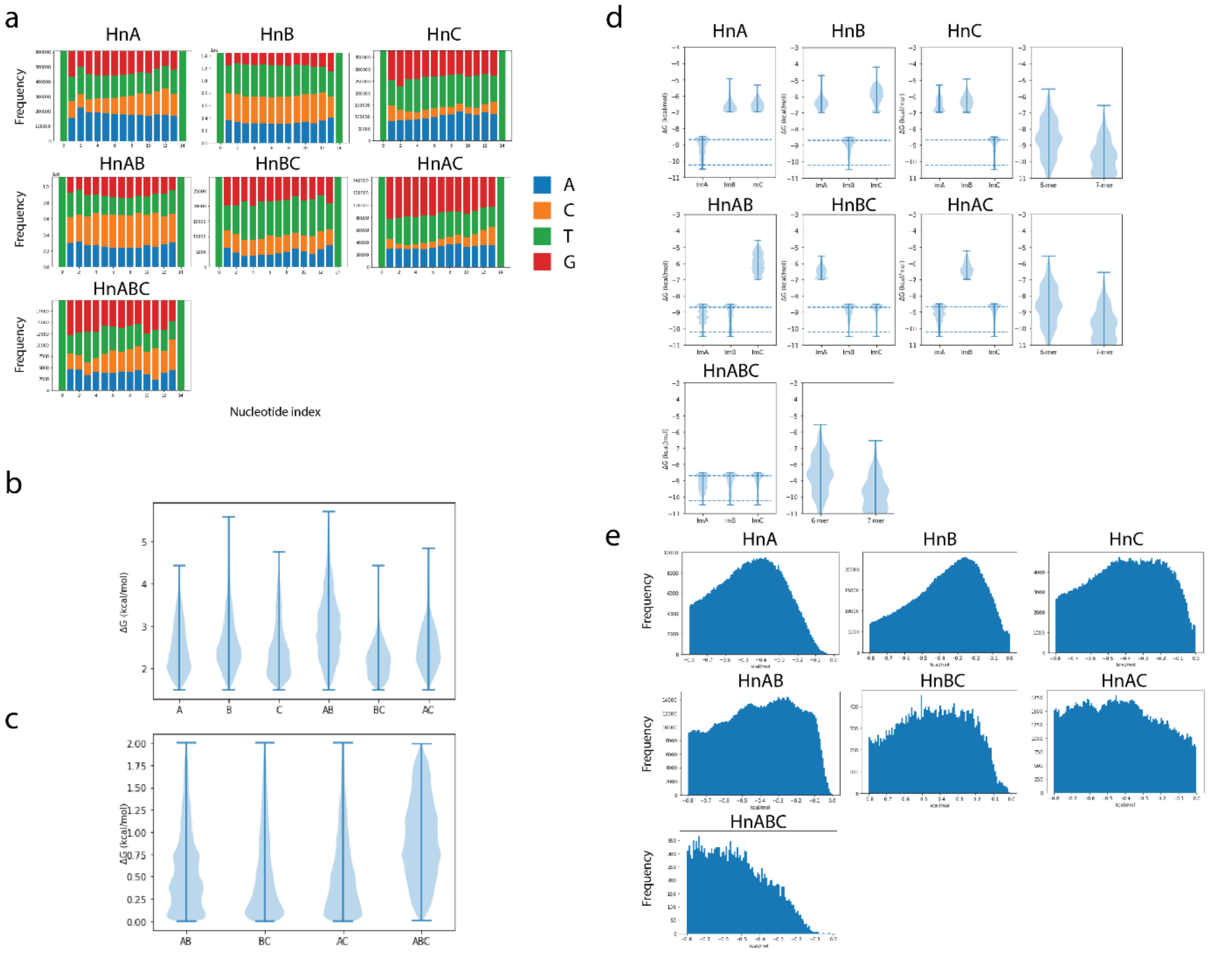

**Supplementary Fig. S2 Parameters of mismatching handles pre-filtering.**

Relevant parameters of mismatching handles coming from the exhaustive search before filtering. **a**, Base preference for each handle species by nucleotide index. From top to bottom: A (blue), C (orange), T (green), G (red). **b**, Wanted-unwanted binding energy gap (difference between the highest energy (weakest binding) among desired bindings and the lowest energy (strongest binding) among undesired binding). **c**, Binding energy discrepancy (maximum difference between the highest and lowest energies of the desired bindings). **d**, Energy of binding between the handles and the imagers. On the right is shown the violin plot of the energy of binding of all possible 6- and 7-mers to their reverse complements in the same conditions as for the mismatching strand, and the dashed lines correspond to their averages. This is to give a comparison between the binding energies of the mismatching sequences and the ones of speed-optimized DNA-PAINT. **e**, Energy of self-interaction.

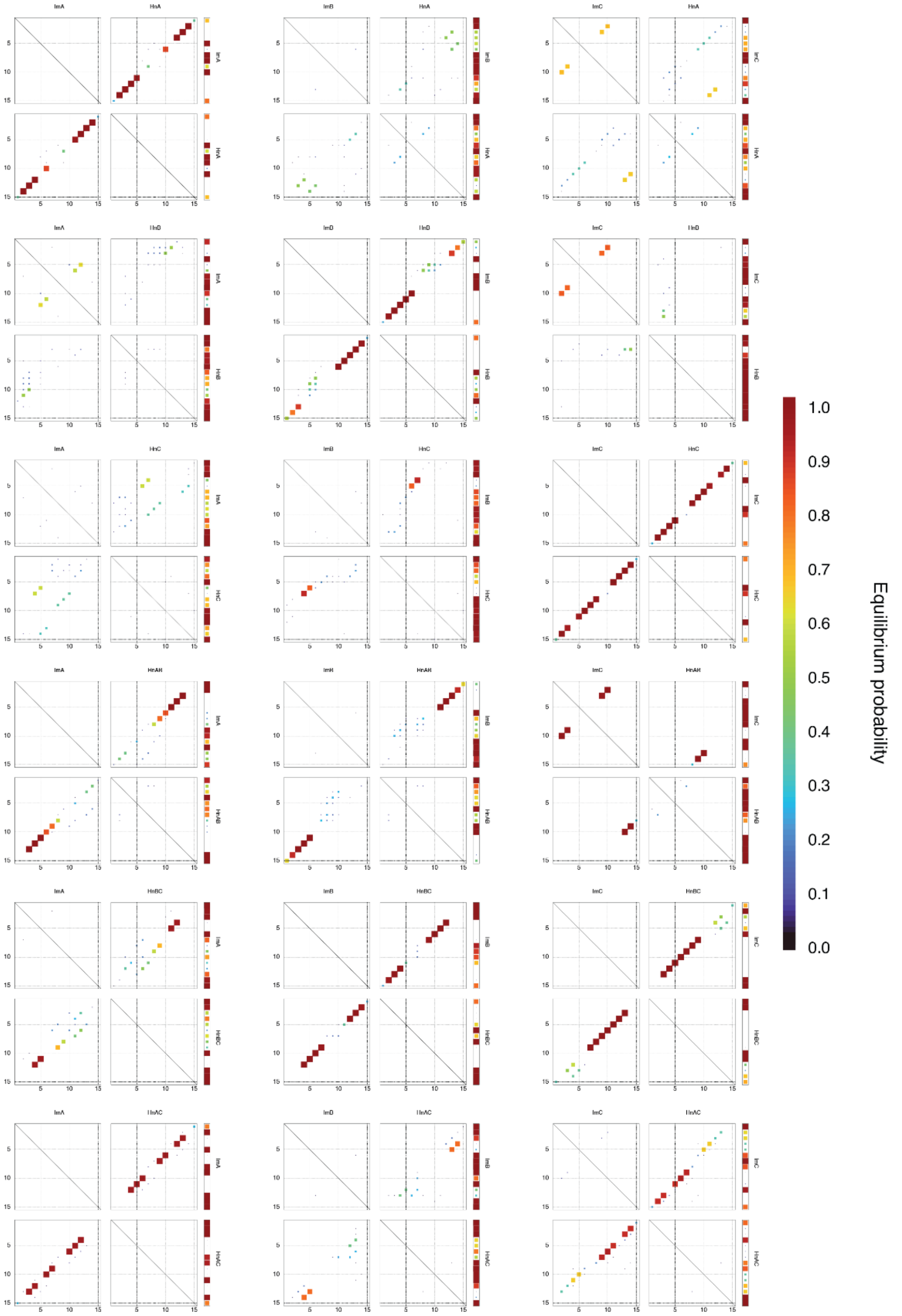

**Supplementary Fig. S3 NUPACK predictions of the selected mismatching strands.**

Pair probabilities of imager strands and handles, as predicted by NUPACK. Predictions were calculated at 0°C, 0.5 M NaCl.

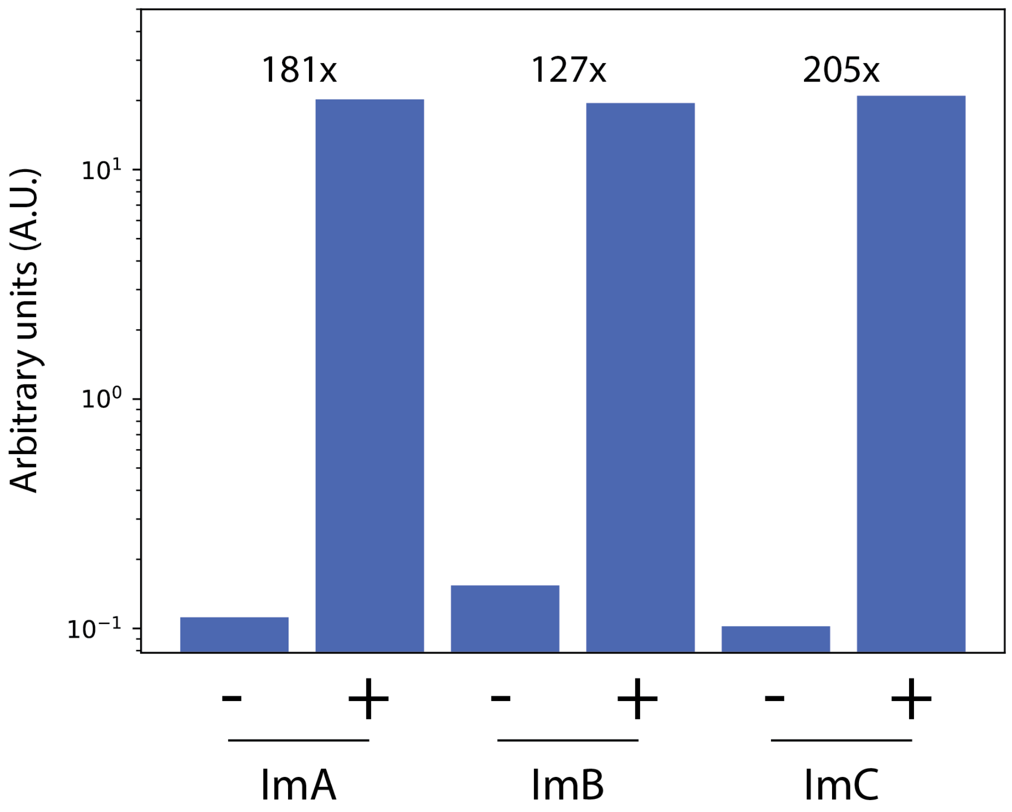

**Supplementary Fig. S4 Fluorogenicity factors of green fluorogenic imagers.**

Fluorogenicity factors for the Cy3b-ImA/B/C-BHQ2 strands, measured as described in Methods. The -/+ refers to the imager in the absence/presence of its reverse complement. Fluorogenicity factors are on top.

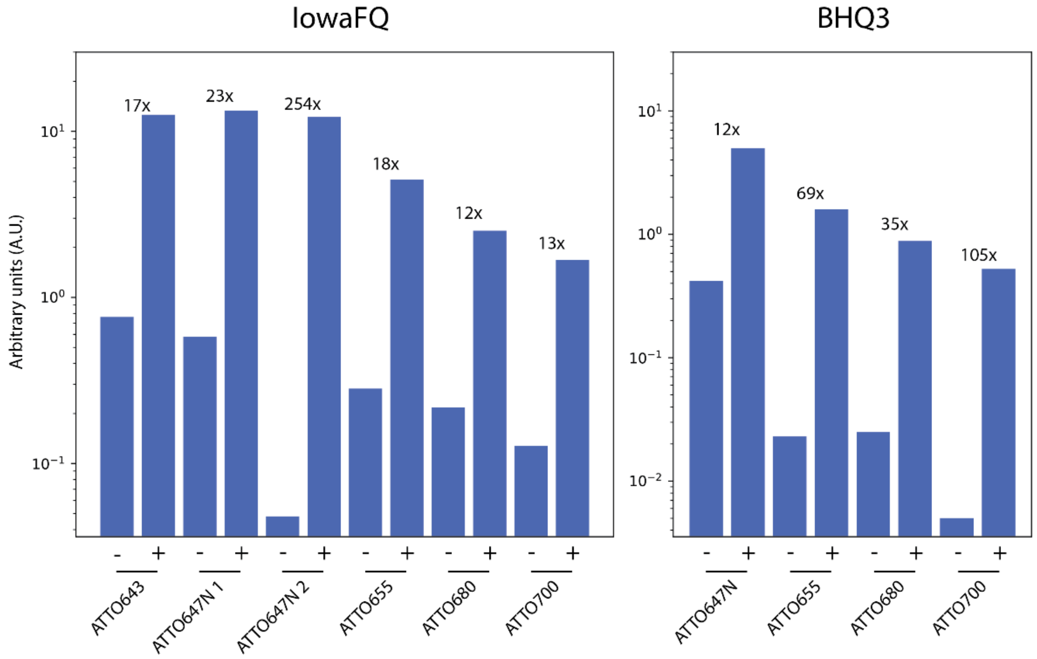

**Supplementary Fig. S5 Fluorogenicity factors of far-red fluorogenic strands.**

Fluorogenicity factors for the far-red ImB strands, measured as described in Methods. The -/+ refers to the imager in the absence/presence of its reverse complement. Fluorogenicity factors. Two quenchers were tested, IowaFQ and BHQ3. Fluorogenicity factors are on top. While BHQ3 showed higher fluorogenicity ratios on average, it also seemed to quench the dye in the bound state more than IowaFQ, leading to lower brightness in the bound state. If the setup is not limited by laser power, however, the higher fluorogenicity factor of BHQ3-labelled imagers could be of interest. Also, ATTO647N 1 and ATTO647N 2 refer to two different HPLC fractions of the ATTO647N conjugation. While ATTO647N 1 exhibits a fluorogenicity factor in line with the other dyes, ATTO647N 2 shows an exceptionally high factor, which could be of interest if ATTO647N is a suitable dye for an application.

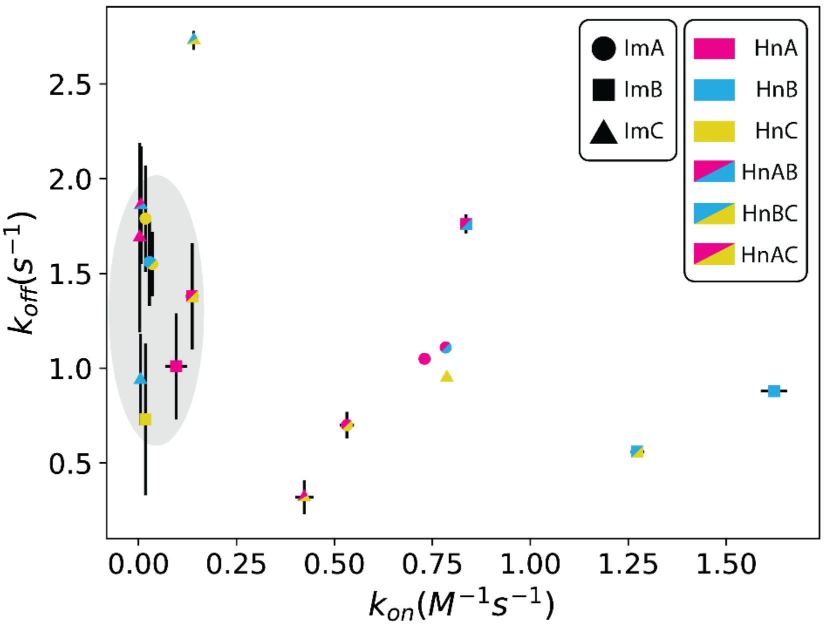

**Supplementary Fig. S6 Kinetics of mismatching handles.**

Average kinetics ± s.e.m of the handles-imager pairs. S.e.m. bars might be hidden by the marker. Undesired bindings are highlighted by the gray ellipse.

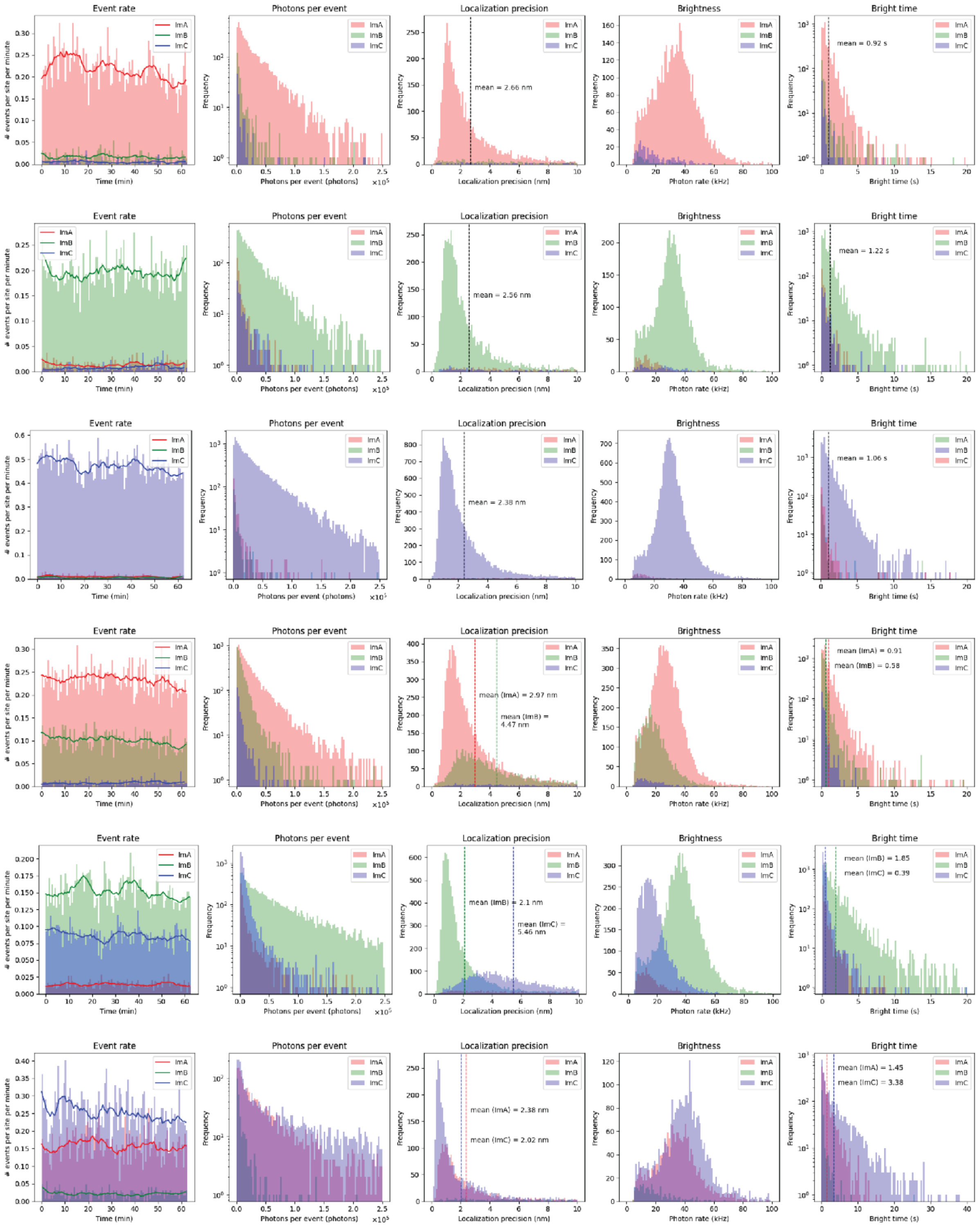

**Supplementary Fig. S7 Photophysics of fluorogenic imagers/handles family.**

Imaging statistics for the chosen mismatching handles.

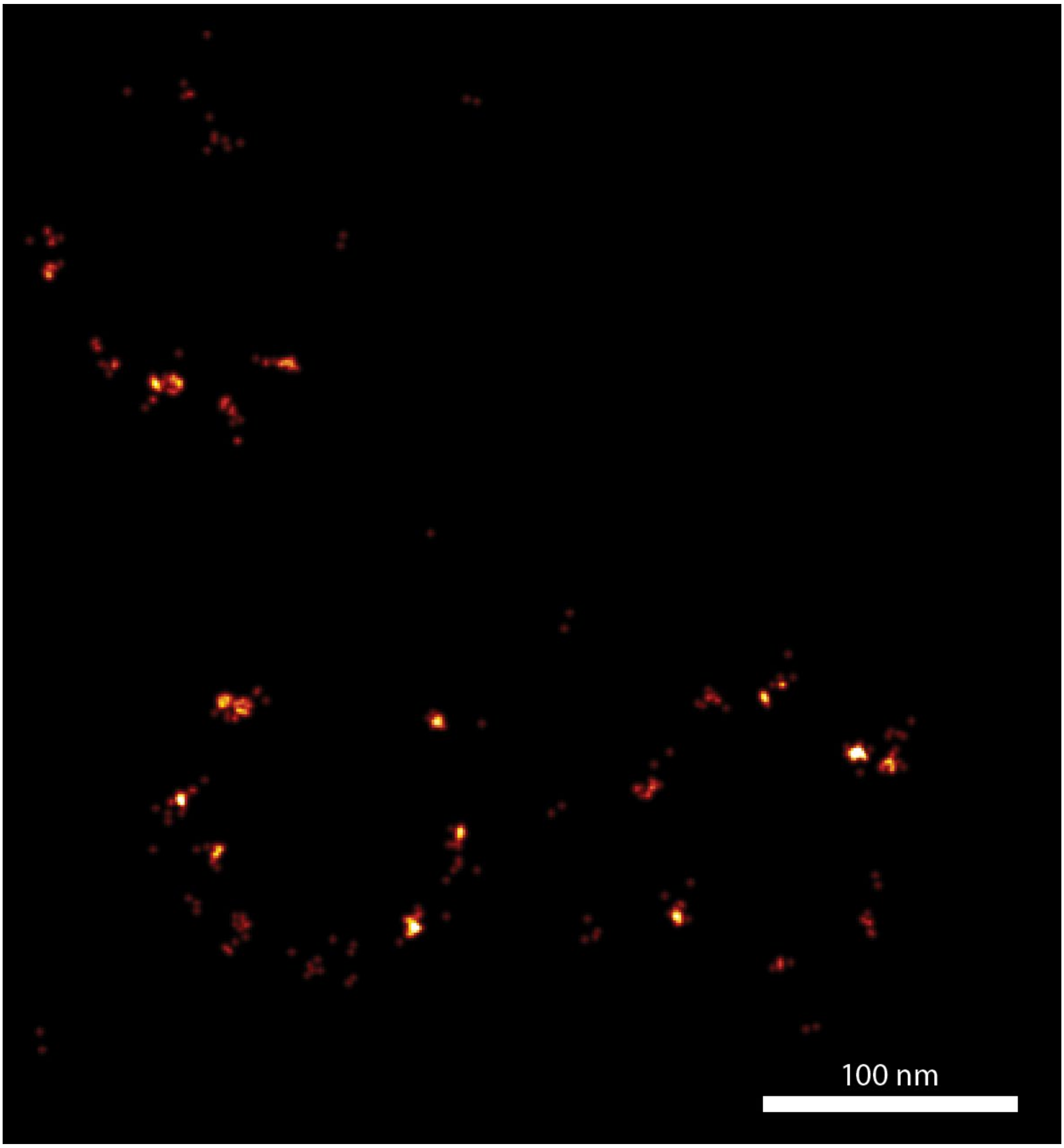

**Supplementary Fig. S8 Nuclear pore complex imaging with mismatching imager.**

2D MINFLUX imaging of a CRIPSR U2OS Nucleoporin96-GFP cell line, labelled with an αGFP nanobody tagged with HnA. The imager was acquired over the course of 20 minutes, with 2.5 nM of the fluorogenic imager ATTO655-ImA-IowaFQ.

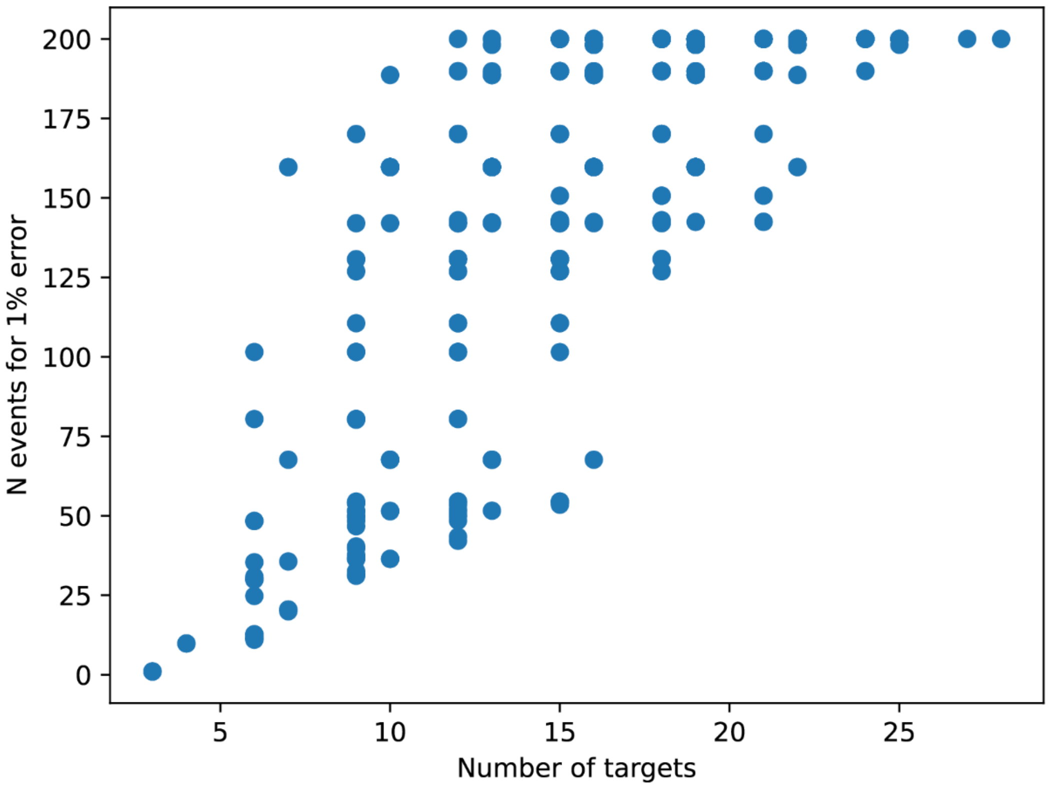

**Supplementary Fig. S9 Efficiency of modular multiplexing stoichiometries.**

Scatter plot of the efficiency (as in number of events required to reach a 1% error) across different number of targets and different stoichiometries of binding sites on the modular multiplexing binding site. Even for comparatively low number of targets (e.g., 6), a large range of efficiencies can be achieved, making the choice of stoichiometries a non-trivial and important problem. The required number of events was clipped at 200 if higher.

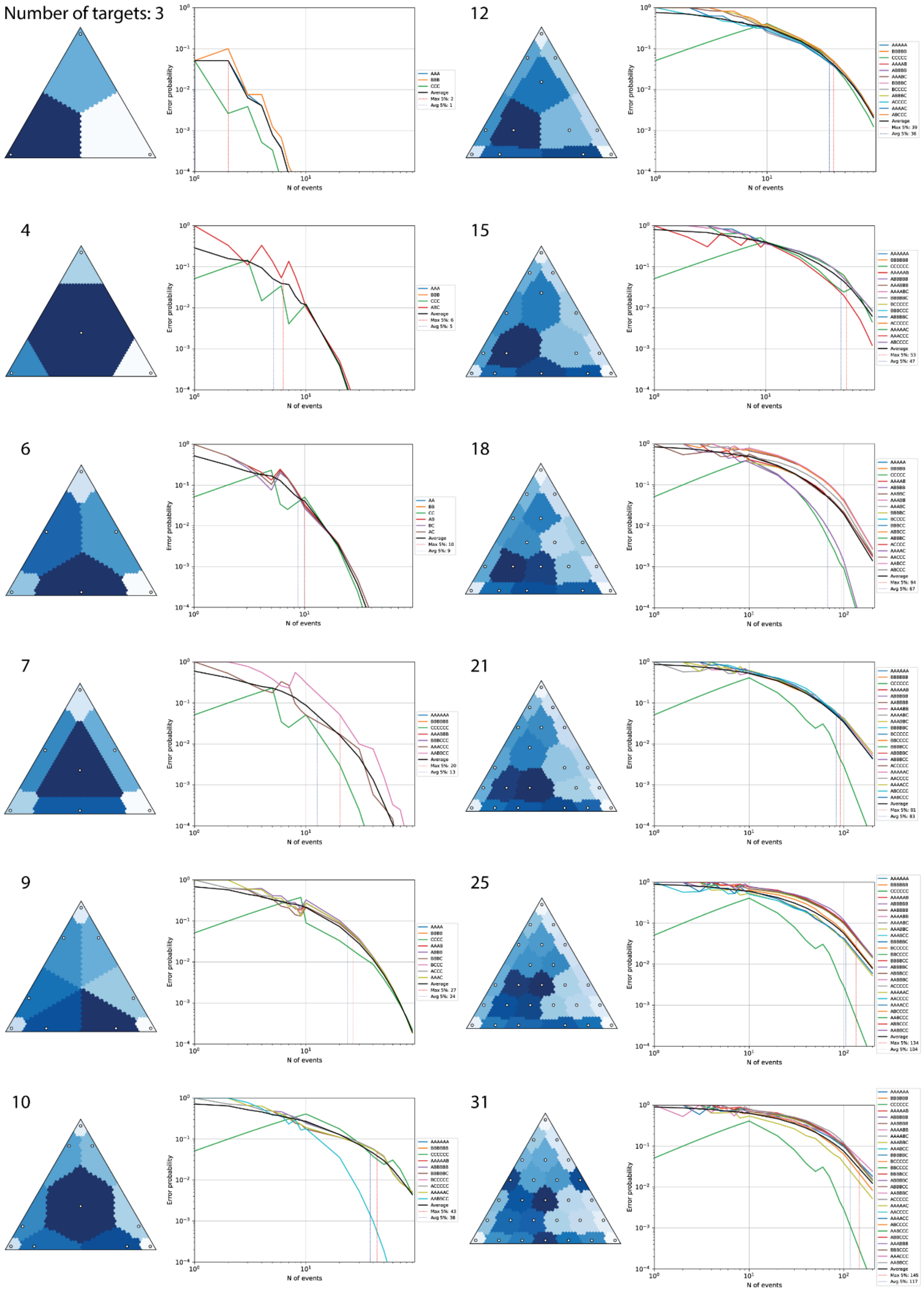

z

**Supplementary Fig. S10 Optimal module stoichiometries and error probabilities.**

Optimal stoichiometries and associated error probabilities versus number of events, by number of targets. All combinations of 3, 4, 5 and 6 modules were simulated, and the most efficient stoichiometry to distinguish a number of targets from 3 to 28 was chosen based on the number of events required for the error probability to cross a 5% threshold. The ternary plots on the left side show the strategy, with the maximum likelihood regions of each species in different colors. The plots on the right side show the error probability versus the number of events. These simulations were done assuming a ~5% off-target binding, which is in line with the experimental ones (1% < < 10%, for different handles across the modular and mismatching strategies).

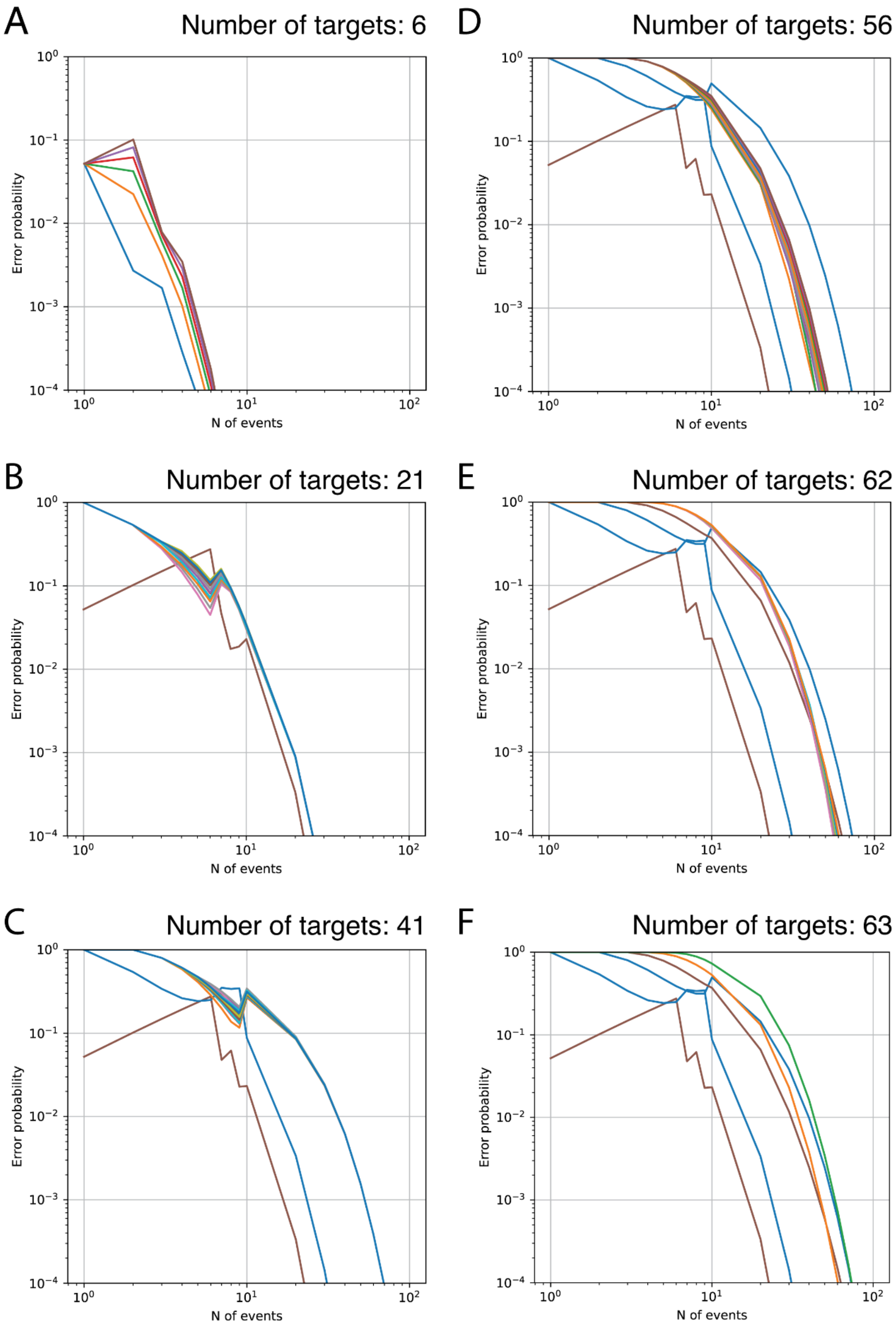

**Supplementary Fig. S11 Error probabilities for a non-stoichiometric, 6-label approach.**

Theoretical performance of a 6-label approach, as opposed to the 3-label approach used in this work. The legends are omitted for lack of enough separation and colors in the plot. All simulations assume a ~5% off-target binding, as in Supplementary Fig. S10 **a**, Simulation of a 6-color approach (where each handle has a probability vector of the form $\mathbf{p} = (p_{R1}, p_{R2}, p_{R3}, p_{R4}, p_{R5}, p_{R6}) = \delta_i$, that is, the vector is 0 at all components except the $i$-th component, at which it is 1). 6 is $\binom{6}{1}$. **b**, Simulation of a 21-color approach ($\mathbf{p} = \delta_i$ or $\frac{1}{2}\delta_{i,j}$, that is, the vector is either as in the 6-color approach, or is 0 at all components except the $i$-th and $j$-th components, at which it is $\frac{1}{2}$, for some $(i,\ j),\ i < j$). 21 is $\binom{6}{1} + \binom{6}{2}$. **c**, Simulation of a 41-color approach ($\mathbf{p} = \delta_i$ or $\frac{1}{2}\delta_{i,j}$ or $\frac{1}{3}\delta_{i,j,k}$, for some $(i,\ j,\ k),\ i < j < k$). 41 is $\binom{6}{1} + \binom{6}{2} + \binom{6}{3}$. **d**, Simulation of a 56-color approach ($\mathbf{p} = \delta_i$ or $\frac{1}{2}\delta_{i,j}$ or $\frac{1}{3}\delta_{i,j,k}$ or $\frac{1}{4}\delta_{i,j,k,l}$, for some $(i,\ j,\ k,\ l),\ i < j < k < l$). 56 is $\binom{6}{1} + \binom{6}{2} + \binom{6}{3} + \binom{6}{4}$. **e**, Simulation of a 62-color approach ($\mathbf{p} = \delta_i$ or $\frac{1}{2}\delta_{i,j}$ or $\frac{1}{3}\delta_{i,j,k}$ or $\frac{1}{4}\delta_{i,j,k,l}$ or $\frac{1}{5}\delta_{i,j,k,l,m}$, for some $(i,\ j,\ k,\ l,\ m),\ i < j < k < l < m$). 62 is $\binom{6}{1} + \binom{6}{2} + \binom{6}{3} + \binom{6}{4} + \binom{6}{5}$. **f**, Simulation of a 63-color approach ($\mathbf{p} = \delta_i$ or $\frac{1}{2}\delta_{i,j}$ or $\frac{1}{3}\delta_{i,j,k}$ or $\frac{1}{4}\delta_{i,j,k,l}$ or $\frac{1}{5}\delta_{i,j,k,l,m}$ or $\frac{1}{6}\mathbf{1}$, for some $(i,\ j,\ k,\ l,\ m),\ i < j < k < l < m$) and $\mathbf{1}$ is the vector of ones. 63 is $\binom{6}{1} + \binom{6}{2} + \binom{6}{3} + \binom{6}{4} + \binom{6}{5} + \binom{6}{6}$.

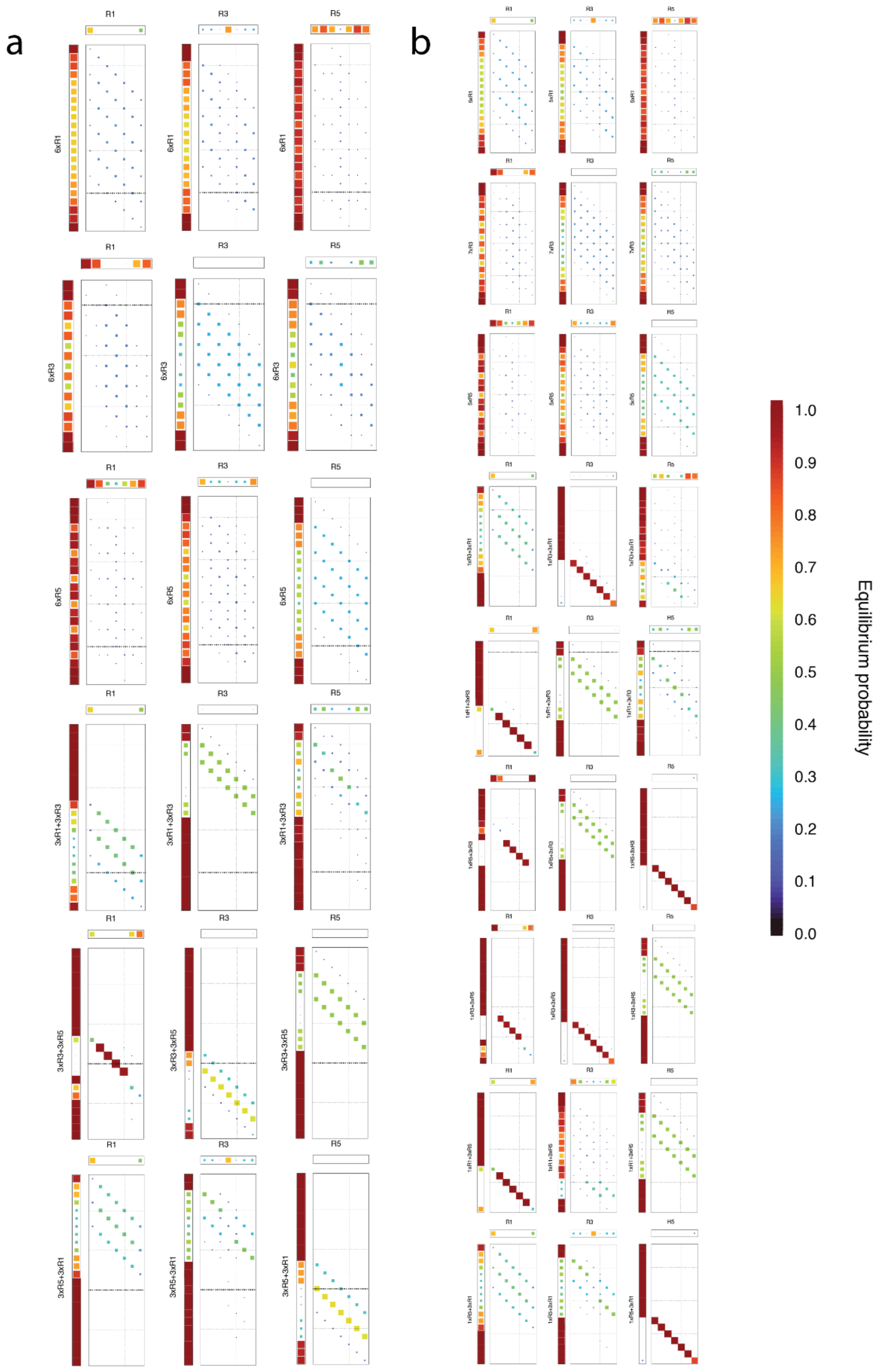

**Supplementary Fig. S12 NUPACK predictions of modular multiplexing.**

Pair probabilities of imager strands and handles for the 6-color (**a**) and 9-color (**b**) multiplexing strategies, as predicted by NUPACK. Predictions were calculated at 0°C, 0.5 M.

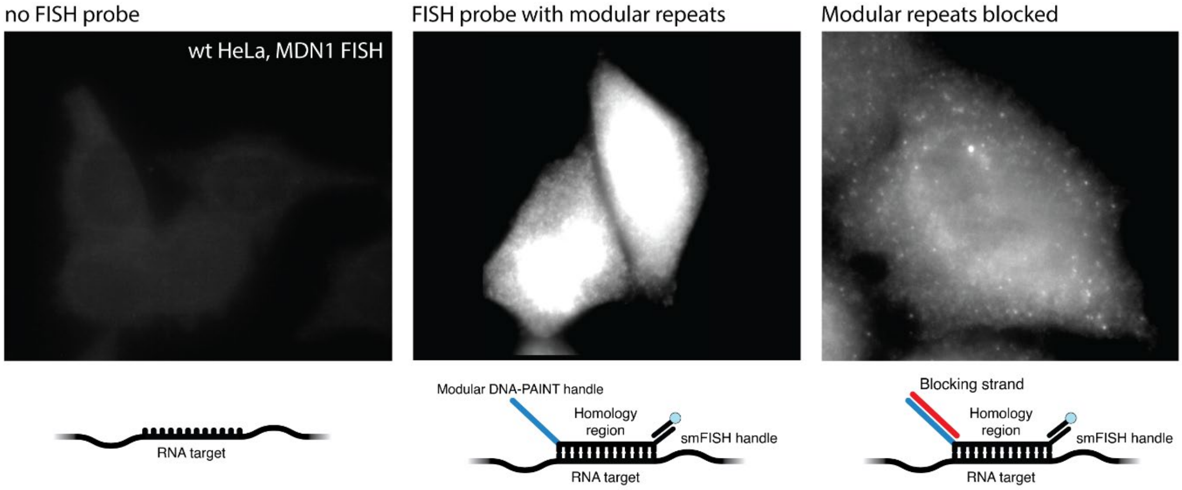

**Supplementary Fig. S13 Module-based probe-induced RNA-FISH background.**

Comparison between the background induced through FISH probes extended with the modular repeat DNA-PAINT handles. Left: negative control of a wt Hela cell with no FISH probes added, Middle: background induced by using the modular repeat extended FISH probes, Right: background is reduced when blocking the modular repeat extension by incubating the FISH probes with a fully complementary DNA strand prior to hybridization. Contrast and brightness are equal across panels.

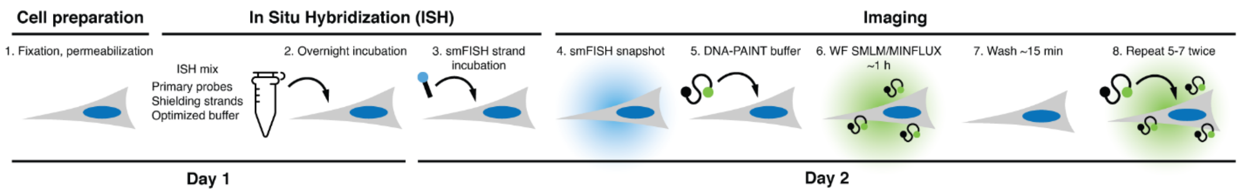

**Supplementary Fig. S14 Graphical protocol of RNA-FISH DNA-PAINT.**

Graphical protocol of RNA-FISH DNA-PAINT. The protocol takes 2 days (without considering cell seeding, which is cell-line dependent). Imaging can be performed immediately, or up to 2 days after sample preparation as described in the Methods. Longer times lead to a decrease in sample labelling efficiency and fluorescence background.

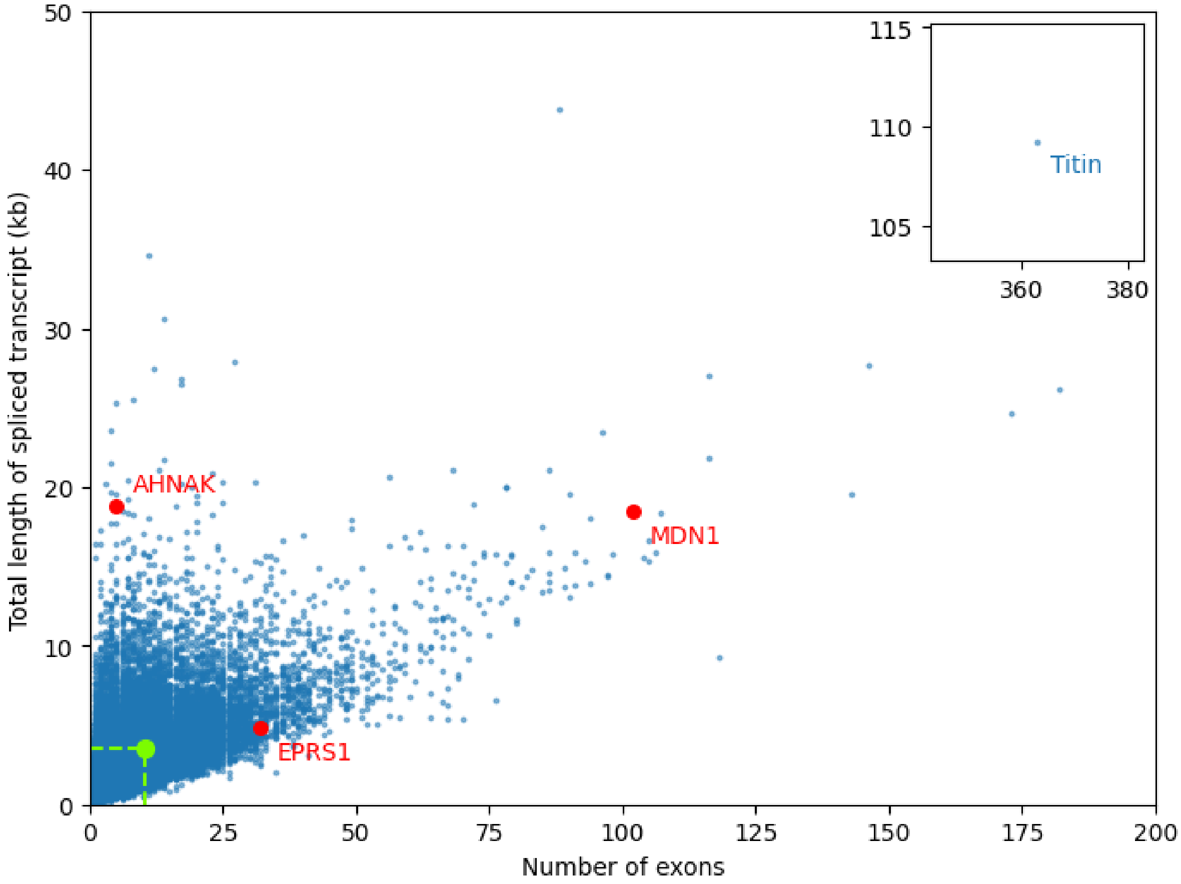

**Supplementary Fig. S15 Gene length versus number of exons in the human genome.**

Overview of number of spliced trascript length versus number of exons for the main isoform (as indicated by the Gencode APPRIS notation of main functional isoform or Transcript Support Level of 1) of all protein-coding human genes. The genes chosen in this study are highlighted, as is the mean of all genes (green).

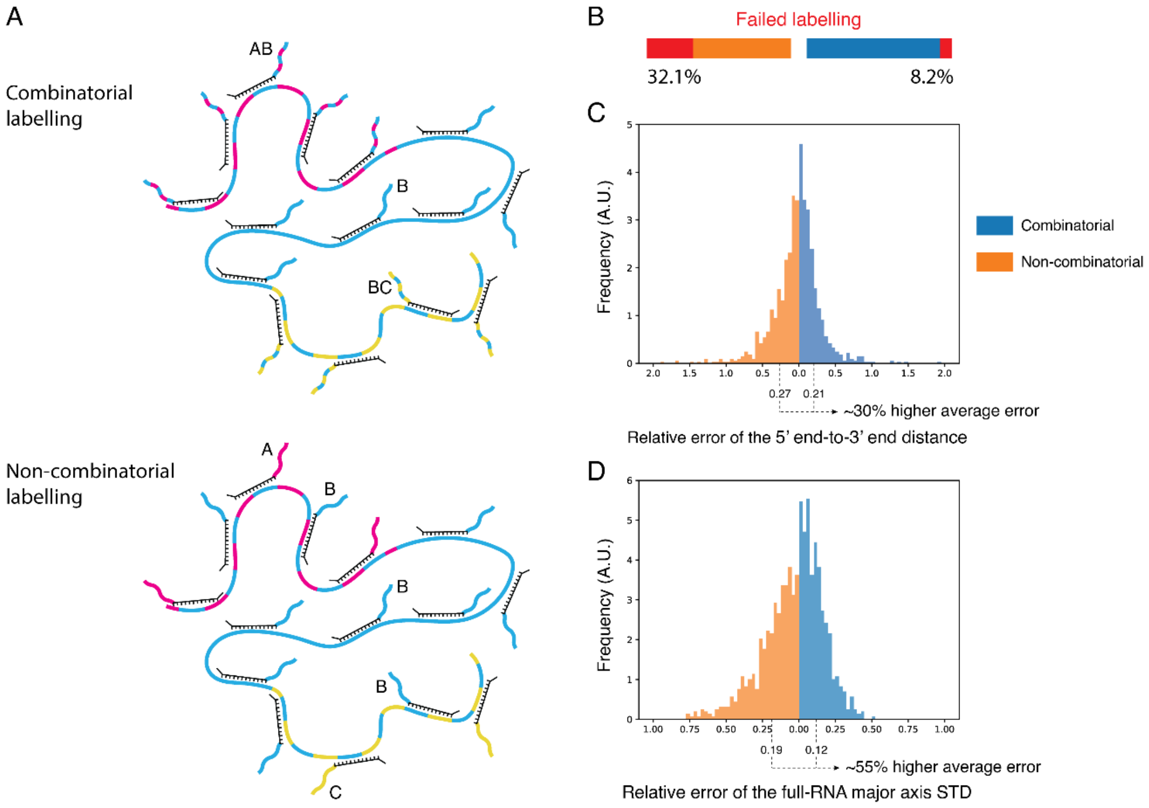

**Supplementary Fig. S16: Combinatorial versus non-combinatorial multiplexing comparison**

Comparison of the performance of a combinatorial multiplexing strategy and non-combinatorial multiplexing strategy. **a**. Top: combinatorial multiplexing strategy analyzed, bottom: non-combinatorial multiplexing strategy analyzed. In each case, the RNA transcript is subdivided into three segment (magenta-cyan, cyan, yellow-cyan). In the combinatorial case, each probe targeting the "combinatorial" regions (magenta-cyan and yellow-cyan) will bind to the magenta imager strand and the cyan imager strand. In the non-combinatorial case, each probe of a "combinatorial" region binds only one type of imager, effectively halving the number of probes for each imager strand. 1000 RNAs were simulated (random walk), each 5 kb long and with 100 probes equi-spaced along its length, assuming a 20% labelling efficiency and an average of 15 localizations with 5 nm sigma stemming from each probe. **b**. In the non-combinatorial case, the reduced number of probes combined with the stochasticity of probe binding increases the probability of not detecting any magenta or yellow signal by 4-fold. **c-d**. The under-labelling also affects the metrics reported from the transcript like 5' end-to-3' end distance (by about 30%, **c**) or major axis standard deviation of the entire RNA (cyan signal, by about 50%, **d**. Relative error $= |\mathrm{metric}_{\mathrm{combi\ or\ non-combi}} - \mathrm{metric}_{\mathrm{true}}| / \mathrm{metric}_{\mathrm{true}}$, where $\mathrm{metric}_{\mathrm{combi\ or\ non-combi}}$ refers to the value of the metric (5' end-to-3' end distance or major axis standard deviation) extracted from the simulated localizations in the combi or the non-combi case, while $\mathrm{metric}_{\mathrm{true}}$ refers to the true value of that metric in that simulation. The results presented hold true for any combination of simulation parameters, however, as they are a consequence of the under-sampling of non-combinatorial labelling.

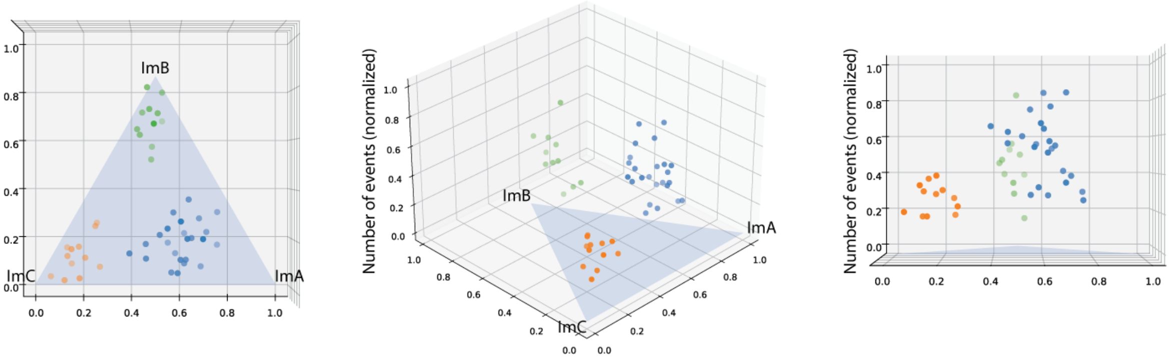

**Supplementary Fig. S17 RNA-FISH Combi-PAINT with event number information.**

RNA-FISH Combi-PAINT blind classification can be improved further by including absolute number of events information, due to the mRNAs being labelled with different number of probes due to length and/or sequence-specificity constraints. The total number of events from all rounds was normalized to allow comparison with the proportion of binding to each imager strand.

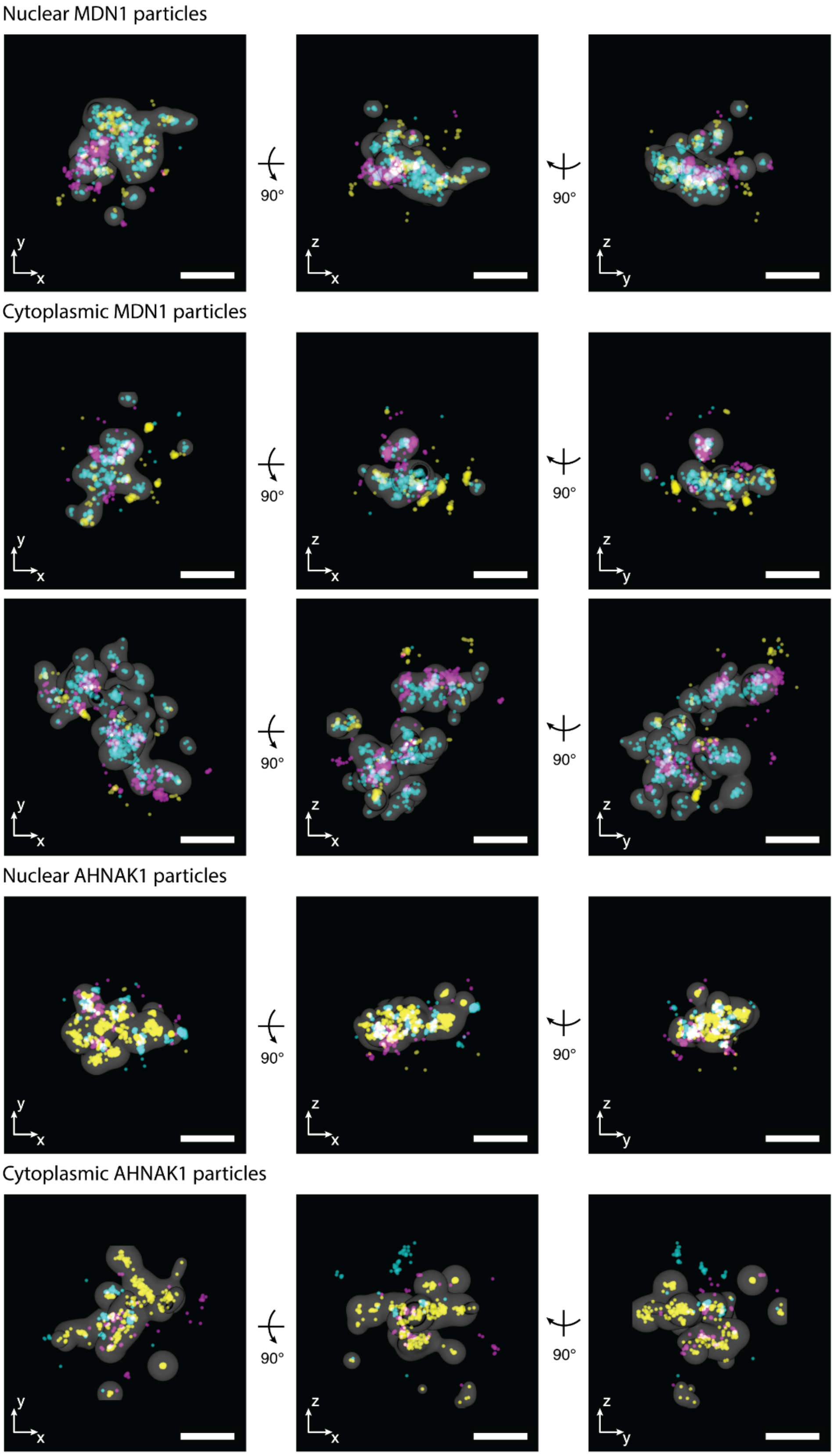

**Supplementary Fig. S18 Additional 3D MINFLUX MDN1 and AHNAK1 mRNPs**

Additional MDN1 and AHNAK1 3D MINFLUX mRNPs, marked as nuclear or cytoplasmic based on their localization

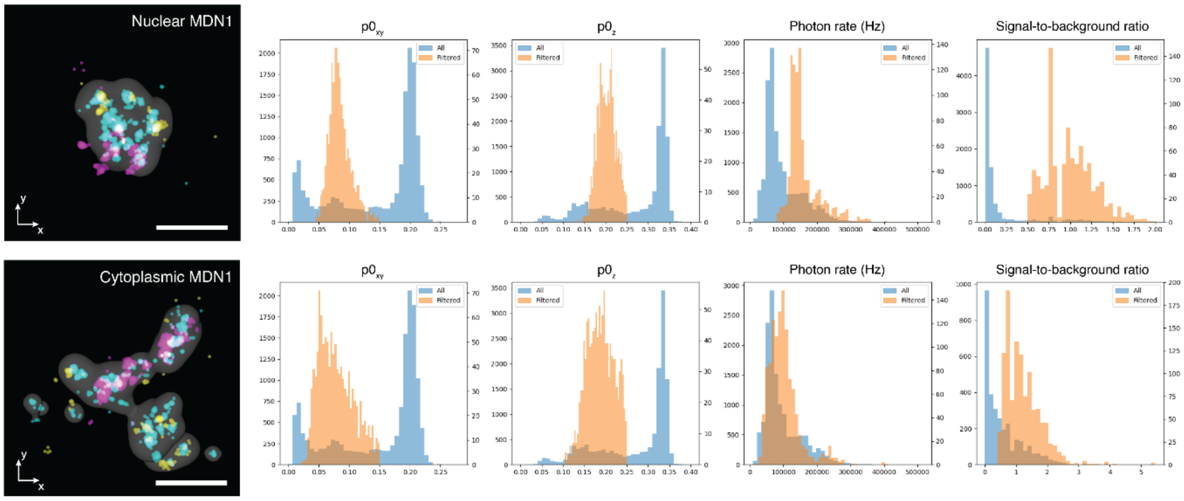

**Supplementary Fig. S19 3D MINFLUX filtering parameters**

Typical parameters and thresholds used for the filtering of 3D MINFLUX localizations. $p_{0\ xy}(n_{0,vortex}/N_{vortex},$ the ratio of the photons from the central vortex in the last iteration and all vortex exposures in the last iteration), $p_{0\ z}(n_{0,top\ hat}/N_{top\ hat},$ the ratio of the photons from the central top hat in the last iteration and all top hat exposures in the last iteration) and SBR (signal-to-background ratio). The photon rate (in Hz) can be used to filter double-blinking events, which are however also addressed by the $p_0$ ratios.

#### Supplementary Tables

| Handle species | Matches with ImA | Matches with ImB | Matches with ImC | End matches | ΔG discrepancy (kcal/mol) | Wanted/ unwanted ΔG gap (kcal/mol) | ΔG self (kcal/mol) | ΔG ImA (kcal/mol) | ΔGImB (kcal/mol) | ΔG ImC (kcal/mol) |
|---|---|---|---|---|---|---|---|---|---|---|
| HnA | > 9 | < 8 | < 8 | > 2 | / | > 2 | > -0.3 | < -9.5 | / | / |
| HnB | > 9 | < 7 | < 7 | > 2 | / | > 2 | > -0.3 | / | < -9.2 | / |
| HnC | > 9 | < 7 | < 7 | > 2 | / | > 2 | > -0.3 | / | / | < -9.2 |
| HnAB | > 9 | > 9 | / | / | < 0.5 | > 2 | > -0.3 | < -9.2 | < -9.2 | / |
| HnBC | / | > 8 | > 8 | / | < 0.5 | > 2 | > -0.4 | / | < -9.2 | < -9.2 |
| HnAC | > 8 | < 8 | > 8 | / | < 0.5 | > 2 | > -0.4 | < -9.2 | / | < -9.2 |
| HnABC | > 5 | > 5 | > 5 | / | < 0.6 | / | > -0.6 | < -9.2 | < -9.2 | < -9.2 |

**Supplementary Table S1 Filtering parameters for mismatching handles screening.**

List of filters used during mismatching handle screening. “Matches with ImA/B/C” refers to matches between the imager and handle sequences aligned in frame (no secondary structure considered). “End matches” refers to a minumum number of consecutive matches at the 5’ and 3’. “ΔG discrepancy” refers to the maximum difference between the highest and lowest energies of the desired bindings (hence the handle must bind at least two imagers for this value to be available). “Wanted/unwanted ΔG gap” refers to the minimum difference between the highest energy (weakest binding) among desired bindings and the lowest energy (strongest binding) among undesired binding (hence the handle must not bind at least one imager for this value to be available). “ΔG self” refers to the energy of self-interaction, while “ΔG ImA/B/C” refers to the energy of binding between the handle and the imager.

| Experiment | Imaging parameters | Imaging buffer | Microscope |
|---|---|---|---|
| Initial imager screening | 20 ms, 100k frames, 45 $W/cm^2$ | ImA- or ImB- or ImC-Cy3b: 10 nM, B+PCA/PCD/trolox, 12.5 mM $MgCl_2$ | Elyra 7 |
| Figure 2, SI Figure 4-5 (6 color mismatching) | 150 ms, 25k/25k/25k frames, 380 $W/cm^2$ | ImA/B/C-Cy3b: 5/2/10 nM, B+PCA/PCD/trolox, 12.5 mM $MgCl_2$ | Custom setup, PCO edge 26 |
| Figure 2 (6 col modular) | 100 ms, 25k/25k/30k frames, 380 $W/cm^2$ | R1/3/5-Cy3b: 500/750/1500 pM, B+PCA/PCD/trolox, 12.5 mM $MgCl_2$ | Custom setup, PCO edge 26 |
| Figure 2 (9 col modular) | 100 ms, 25k/25k/35k frames, 380 $W/cm^2$ | R1/3/5-Cy3b: 320/480/640 pM, B+PCA/PCD/trolox, 50 mM $MgCl_2$ | Custom setup, PCO panda 4.2 bi |
| Widefield cell experiments | 150 ms, 25k frames per round, ~400 $W/cm^2$ | ImA/B/C-Cy3b: 10/5/15 nM, 1xPBS + 500 mM NaCl + PCA/PCD/trolox | Elyra 7 |

**Supplementary Table S2 Imaging conditions of the widefield experiments.**

Experimental conditions for widefield experiments. For the screening, each origami was measured with only one strand in solution, meaning for the mixed handles (HnAB, HnBC, HnAC) two separate experiments were done. The other experiments are Exchange PAINT experiments done over 3 rounds of imaging (R1/ImA -> R3/ImB -> R5/ImC) in the respective conditions.

| Handle name | Notes | Sequence (5'->3'), including linker and padding |
|---|---|---|
| HnA | 6 color mismatching, HnA8 in the screen | A **TATGTTACAAGGCCT** CG |
| HnB | 6 color mismatching, HnB7 in the screen | T **TAGTCTCCCCCTCAT** CC |
| HnC | 6 color mismatching, HnC7 in the screen | T **TGGTTCGATAATACT** AG |
| HnAB | 6 color mismatching, HnAB3 in the screen | T **TATTTATTGGCGCAT** TT |
| HnBC | 6 color mismatching, HnBC5 in the screen | T **TAGTTGTATCCGAGT** TT |
| HnAC | 6 color mismatching, HnAC6 in the screen | T **TGGGTGTGGAGGCGT** AT |
| 6xR1 | 6 color modular | TC **TCCTCCT**CCTCCTCCTCCTCCT |
| 6xR3 | 6 color modular | TC **CTCTCTC**TCTCTCTCTC |
| 6xR5 | 6 color modular | CC **CTTCTTC**TTCTTCTTCTTCTTC |
| 3xR1+3xR3 | 6 color modular | TC **TCCTCCT**CCTCCTCT**CTCTCTC** |
| 3xR3+3xR5 | 6 color modular | TC **CTCTCTC**TCTCCTTCTT**CTTCTT C** |
| 3xR5+3xR1 | 6 color modular | CC **CTTCTTC**TTCTTCCTCC**TCCTCCT** |
| 5xR1 | 9 color modular | TC **TCCTCCT**CCTCCTCCTCCT |
| 7xR3 | 9 color modular | TC **CTCTCTC**TCTCTCTCTCTC |
| 5xR5 | 9 color modular | CC **CTTCTTCTTCTTCTTCTTC** |
| 1xR3+3xR1 | 9 color modular | TC **CTCTCTC**CTCC**TCCTCCT** |
| 1xR1+3xR3 | 9 color modular | TC **TCCTCCT**CT**CTCTCTC** |
| 1xR5+3xR3 | 9 color modular | CC **CTTCTTC**CCTCTCT**CTCTCTC** |
| 1xR3+3xR5 | 9 color modular | TC **CTCTCTC**CCTTCTT**CTTCTTC** |
| 1xR1+3xR5 | 9 color modular | TC **TCCTCCT**TCTT**CTTCTTC** |
| 1xR5+3xR1 | 9 color modular | CC **CTTCTTC**CTCC**TCCTCCT** |

**Supplementary Table S3 List of handle sequences.**

List of the handles (mismatching and modular) used in this work. Each handle includes a dinucleotide linker (5' for modular sequences, 3' for mismatching sequences) and mismatching sequences have a 5' extra nucleotide as padding as in (*39*). Both linked and padding nucleotides were computed in NUPACK to minimize changes in the energies of self-interaction (for mismatching handles only) and of interaction with the imagers (for both handle types). Binding motifs are bold and underlined. For modular handles, the corresponding motifs are colored and shared nucleotides are in black; spacing nucleotides are highlighted in gray.

| | ImA | | ImB | | ImC | | GC fraction (%) |
|---|---|---|---|---|---|---|---|
| | $\Delta G_{binding}$ (kcal/mol) | Mismatches | $\Delta G_{binding}$ (kcal/mol) | Mismatches | $\Delta G_{binding}$ (kcal/mol) | Mismatches | |
| **HnA** | -9.65 | 4 | -6.26 | 9 | -6.95 | 9 | 40.00 |
| **HnB** | -5.89 | 10 | -9.63 | 4 | -5.48 | 11 | 53.33 |
| **HnC** | -6.58 | 10 | -6.25 | 10 | -9.85 | 3 | 33.33 |
| **HnAB** | -9.72 | 5 | -9.98 | 5 | -6.86 | 9 | 33.33 |
| **HnBC** | -6.57 | 8 | -10.24 | 5 | -10.47 | 4 | 40.00 |
| **HnAC** | -10.14 | 6 | -6.12 | 9 | -9.72 | 6 | 66.66 |
| GC fraction (%) | 46.67 | | 40 | | 33.33 | | |

**Supplementary Table S4 List of thermodynamic parameters.**

List of thermodynamic parameters for the imager-handle pairs presented. Each imager-handle combination reports the predicted binding energy in kcal/mol (calculated with NUPACK, calculated for 25°C, 50 mM $Na^+$ and 10 mM $Mg^{2+}$ with model “dna04-nupack3” and ensemble “some-nupack3”) and number of mismatches for the aligned sequences. The last row/column contains the % GC content of the sequence in the column/row.

| Product | 5' mod. | Oligo sequence | 3' mod. | Supplier | Conjugate |
|---|---|---|---|---|---|
| Cy3b-ImA-BHQ2 | Amine | AGGCGTCATCACATA | BHQ2 | Biosearch Technologies | Cy3b-NHS ester |
| Cy3b-ImB-BHQ2 | Amine | ATGCGGAAAAGACTA | BHQ2 | Biosearch Technologies | Cy3b-NHS ester |
| Cy3b-ImC-BHQ2 | Amine | AGTCTTATACAACCA | BHQ2 | Biosearch Technologies | Cy3b-NHS ester |
| ATTO643-ImA- IWFQ | Amine | AGGCGTCATCACATA | IWFQ | IDT | ATTO643-maleimide |
| ATTO643-ImB-IWFQ | Thiol | ATGCGGAAAAGACTA | IWFQ | IDT | ATTO643-maleimide |
| ATTO643-ImC- IWFQ | Amine | AGTCTTATACAACCA | IWFQ | IDT | ATTO643-maleimide |
| ATTO647N-ImB-IWFQ | Thiol | ATGCGGAAAAGACTA | IWFQ | IDT | ATTO647N-maleimide |
| ATTO655-ImA- IWFQ | Thiol | AGGCGTCATCACATA | IWFQ | IDT | ATTO655-maleimide |
| ATTO655-ImB- IWFQ | Thiol | ATGCGGAAAAGACTA | IWFQ | IDT | ATTO655-maleimide |
| ATTO680-ImB-IWFQ | Thiol | ATGCGGAAAAGACTA | IWFQ | IDT | ATTO680-maleimide |
| ATTO700-ImB-IWFQ | Thiol | ATGCGGAAAAGACTA | IWFQ | IDT | ATTO700-maleimide |
| ATTO647N-ImB-BHQ3 | Thiol | ATGCGGAAAAGACTA | BHQ3 | IDT | ATTO647N-maleimide |
| ATTO655-ImB-BHQ3 | Thiol | ATGCGGAAAAGACTA | BHQ3 | IDT | ATTO655-maleimide |
| ATTO680-ImB-BHQ3 | Thiol | ATGCGGAAAAGACTA | BHQ3 | IDT | ATTO680-maleimide |
| ATTO700-ImB-BHQ3 | Thiol | ATGCGGAAAAGACTA | BHQ3 | IDT | ATTO700-maleimide |
| R1-Cy3b | | AGGAGGA | Amine | Sigma Aldrich | Cy3b-NHS ester |
| R3-Cy3b | | GAGAGAG | Amine | Sigma Aldrich | Cy3b-NHS ester |
| R5-Cy3b | | GAAGAAG | Amine | Sigma Aldrich | Cy3b-NHS ester |

**Supplementary Table S5 List of imager strand conjugations.**

List of all performed conjugations. IWFQ is the Iowa Black FQ quencher.

| | ImA ($10^6$ $M^{-1}$ $s^{-1}$) | ImB ($10^6$ $M^{-1}$ $s^{-1}$) | ImC ($10^6$ $M^{-1}$ $s^{-1}$) |
|---|---|---|---|
| **HnA** | 0.730 ± 0.016 | 0.096 ± 0.028 | 0.003 ± 0.002 |
| **HnB** | 0.028 ± 0.005 | 1.622 ± 0.034 | 0.005 ± 0.001 |
| **HnC** | 0.018 ± 0.004 | 0.018 ± 0.008 | 0.787 ± 0.011 |
| **HnAB** | 0.784 ± 0.011 | 0.836 ± 0.016 | 0.006 ± 0.002 |
| **HnBC** | 0.035 ± 0.006 | 1.273 ± 0.017 | 0.141 ± 0.003 |
| **HnAC** | 0.532 ± 0.018 | 0.137 ± 0.016 | 0.423 ± 0.023 |

**Supplementary Table S6 Association rate constant between fluorogenic imager-handle pairs.**

Association rate constants ($k_{on}$) for the fluorogenic imager-handle family. ±: s.e.m.

| | ImA<br>($s^{-1}$) | ImB<br>($s^{-1}$) | ImC<br>($s^{-1}$) |
|---|---|---|---|
| **HnA** | 1.05 ± 0.03 | 1.01 ± 0.28* | 1.69 ± 0.5* |
| **HnB** | 1.56 ± 0.23* | 0.88 ± 0.03 | 0.94 ± 0.24* |
| **HnC** | 1.79 ± 0.28* | 0.73 ± 0.40* | 0.95 ± 0.02 |
| **HnAB** | 1.11 ± 0.02 | 1.76 ± 0.05 | 1.86 ± 0.31* |
| **HnBC** | 1.55 ± 0.17* | 0.56 ± 0.02 | 2.73 ± 0.05 |
| **HnAC** | 0.70 ± 0.07 | 1.38 ± 0.28* | 0.32 ± 0.09 |

**Supplementary Table S7 Dissociation rate constant between fluorogenic imager-handle pairs.**

List of $k_{off}$ values for the imager-handle pairs presented ± s.e.m. Values marked by (*) have high errors as these combination kave low $k_{on}$ values.
